# Fsr quorum sensing system restricts biofilm growth and activates inflammation in enterococcal infective endocarditis

**DOI:** 10.1101/2025.02.07.636843

**Authors:** Haris Antypas, Verena Schmidtchen, Willy Isao Staiger, Yanhong Li, Rachel Jing Wen Tan, Kenneth Kok Fei Ng, Cheryl Jia Yi Neo, Shalome Meera Radhesh, Frederick Reinhart Tanoto, Ronni Anderson Gonçalves da Silva, Cristina Colomer Winter, Caroline Manzano, Jun Jie Wong, Kevin Pethe, Barbara Hasse, Silvio Daniel Brugger, Siu Ling Wong, Daria Van Tyne, Annelies S. Zinkernagel, Kimberly A. Kline

## Abstract

Infective endocarditis (IE) is a life-threatening biofilm-associated infection, yet the factors driving biofilm formation remain poorly understood. Here, we identified the Fsr quorum sensing (QS) system of *Enterococcus faecalis* as a potent negative regulator of IE pathogenesis. Using microfluidic and *in vivo* models, we show that Fsr is induced in late IE when bacteria become shielded from blood flow. Deleting Fsr altered biofilm metabolism and promoted robust biofilm growth and gentamicin tolerance *in vivo*. Furthermore, Fsr inactivation attenuated inflammation by disrupting IL-1β cleavage and activation via the Fsr-regulated gelatinase (*gelE*), allowing biofilm to grow unchecked by the immune system. Consistent with our pre-clinical findings, analysis of two IE patient cohorts linked naturally occurring Fsr-deficient *E. faecalis* to prolonged bacteremia. Overall, our findings provide insights into the role of QS in biofilm growth, persistence, and immune evasion in enterococcal IE.

## INTRODUCTION

Quorum sensing (QS) is an interbacterial communication system that regulates biofilm formation and other group behaviors through the production, secretion, and sensing of autoinducer signaling molecules ^1^. QS is activated when autoinducers accumulate to a threshold concentration, correlated with increased bacterial population density ^1^, a condition commonly encountered in biofilms within natural habitats. When activated, QS can promote or inhibit gene expression involved in the extracellular matrix polysaccharide and protein synthesis, protease secretion, and eDNA release ^2,3^. This regulation can drive the maturation or dispersal of biofilms ^2^. However, QS is not solely regulated by population density. Mechanical and chemical cues, such as fluid flow, habitat architecture, and nutrient availability, also impact its activation ^4–10^. For instance, fluid flow can disperse autoinducers through advection *in vitro*, thereby preventing QS activation ^4,5^. Conversely, bacteria growing in confined and structurally complex *in vitro* habitats can entrap autoinducers, facilitating QS activation ^4,6–8^. Furthermore, despite QS being widespread across the bacterial kingdom, QS-defective mutants in bacterial species such as *Enterococcus faecalis*, *Staphylococcus epidermidis*, and *Pseudomonas aeruginosa* have been isolated from patients, indicating that QS activation may not always confer an advantage ^11–14^. This genomic variation combined with the diverse mechanical and chemical cues present in the host, raises a central question: How does QS regulate biofilm formation across different host microenvironments, and how does this regulation shape infection outcomes?

During infection, bacteria encounter diverse cues from the host microenvironment, such as heterogeneous fluid flow, nutrient availability, and tissue architecture^15^. One host microenvironment where bacteria encounter all these cues is the heart valves, which direct blood flow through the heart chambers and major arteries ^16^. Despite experiencing one of the highest blood flow rates in the body, heart valves are not exempt from bacterial colonization and infection. Mechanical or inflammatory lesions on the heart valves can render them susceptible to infective endocarditis (IE) ^17^. These lesions trigger the accumulation of platelets and fibrin, leading to thrombus formation at the site of the valvular damage ^18^. Thrombi serve as a substrate for adhesion by Gram-positive bacteria, such as *E. faecalis, S. aureus*, and *Streptococcus spp* ^19–23^. Once bacteria attach to the thrombus, they can proliferate into a biofilm, leading to the infiltration of inflammatory cells and eventually to the formation of a septic vegetation ^24^. The inflammatory response is often insufficient to clear the infection^25–27^. Treatment of IE typically requires a combination of intravenous antibiotic therapy for up to 6 weeks ^28^, but due to the high prevalence of multidrug-resistant bacteria and recalcitrance of biofilms to antibiotics, 50% of patients will also undergo surgery to remove infected tissue and replace the damaged valves ^29^. Despite these treatment approaches, global IE deaths have risen from 28,754 in 1990 to 66,322 in 2019, with age-standardized mortality of IE increasing from 0.73 to 0.87 per 100,000 people over the same period ^30^. This alarming increase highlights the need to better understand how bacteria adapt to the vegetation microenvironment to improve the diagnosis and treatment of IE.

Limited insights into the biofilm-host interplay on cardiac vegetations constrain advances in the diagnosis and treatment of IE. Most studies have focused on initial bacterial adhesion to the thrombus exterior, a microenvironment that differs significantly from the conditions encountered within the vegetation during biofilm maturation. The vegetation exterior is exposed to intense shear stress forces due to blood flow, while the interior is shielded from them. This differential exposure to blood flow may influence QS activity and nutrient availability, depending on bacterial location at the different stages of infection. In this study, we investigated the hypothesis that the heterogeneous microenvironment of IE vegetations critically influences biofilm development and immune evasion in IE caused by *E. faecalis,* the third most common causative agent of this infection ^31,32^. While the virulence factors that facilitate initial adhesion to thrombus in enterococcal IE have been identified ^33–41^, the regulation of biofilm formation and its impact on disease progression remain largely overlooked. Here, we show that fluid flow prevents the induction of the Fsr QS system in *E. faecalis*. However, as vegetations grow and biofilms become shielded from fluid flow, QS is induced. This QS induction restricts biofilm growth and promotes inflammation by upregulating gelatinase, which proteolytically cleaves and activates IL-1β. We also demonstrate that loss of the Fsr QS system, commonly observed in IE clinical isolates ^12,13^, promotes unchecked biofilm growth, diminishes inflammatory response, and increases tolerance to antibiotic treatment. Furthermore, the absence of the *fsr* locus correlates with prolonged bacteremia and high disease severity in individuals with enterococcal IE. Collectively, these findings provide new insights into the role of QS in biofilm growth, antibiotic tolerance, and immune evasion.

## RESULTS

### Fluid flow prevents Fsr QS system induction via advection

Vegetations in IE present a heterogeneous microenvironment, where the exterior is exposed to high shear stress and advection from blood flow, while the interior remains shielded. At the onset of IE, bacteria in the bloodstream attach to the vegetation exterior and become directly exposed to blood flow. We therefore hypothesized that blood flow might trigger physiological adaptations in *E. faecalis* that favor the establishment of IE. To identify these adaptations, we exposed surface-attached *E. faecalis* OG1RF to a pulsatile flow of media for 30 minutes in a microfluidic system. This flow we applied generated a shear stress of 20 dynes/cm^2^, simulating the shear stress on aortic valve leaflets ^42–44^. Compared to surface-attached bacteria that were incubated without flow, transcriptomic analysis revealed 59 genes upregulated and 167 genes downregulated under flow (log_2_FC > 1.0, FDR < 0.05) (**Fig. 1A, Supplementary File 1**). Gene ontology enrichment analysis revealed an overrepresentation of genes linked to ATPase activity, glycolytic processes, and oxidative stress (**Fig. 1B**), suggesting that *E. faecalis* adapts its metabolism to meet energy demands and adapt to different oxygen levels under flow. Notably, transcription of the *fsr* locus (*fsrABDC*) and its associated regulon (*gelE, sprE, entV, RS04585*) ^41,45,46^ were significantly decreased in the presence of fluid flow (**Fig. 1A**). The *fsr* locus encodes the Fsr quorum sensing (QS) system, which is activated when the extracellular concentration of the autoinducer peptide (AIP) gelatinase biosynthesis-activating pheromone (GBAP) exceeds a critical concentration and is therefore typically activated in direct proportion with bacterial population density ^47^. Given the role of GBAP concentration in Fsr activation, we investigated whether *fsr* downregulation under fluid flow was attributed to GBAP advection. To distinguish between advection- and shear stress-mediated transcriptional regulation, we exposed surface-attached bacteria to low (1 dynes/cm^2^), intermediate (10 dynes/cm^2^), and high shear stress (20 dynes/cm^2^). Quantitative PCR showed that transcription of *fsr*-associated genes was decreased after 30 min, regardless of the shear stress magnitude applied (**Fig. 1C**), demonstrating that advection and not mechanical forces regulate their expression. Taken together these data show that fluid flow impacts quorum sensing and triggers physiological adaptations in *E. faecalis*.

**Fig. 1.**
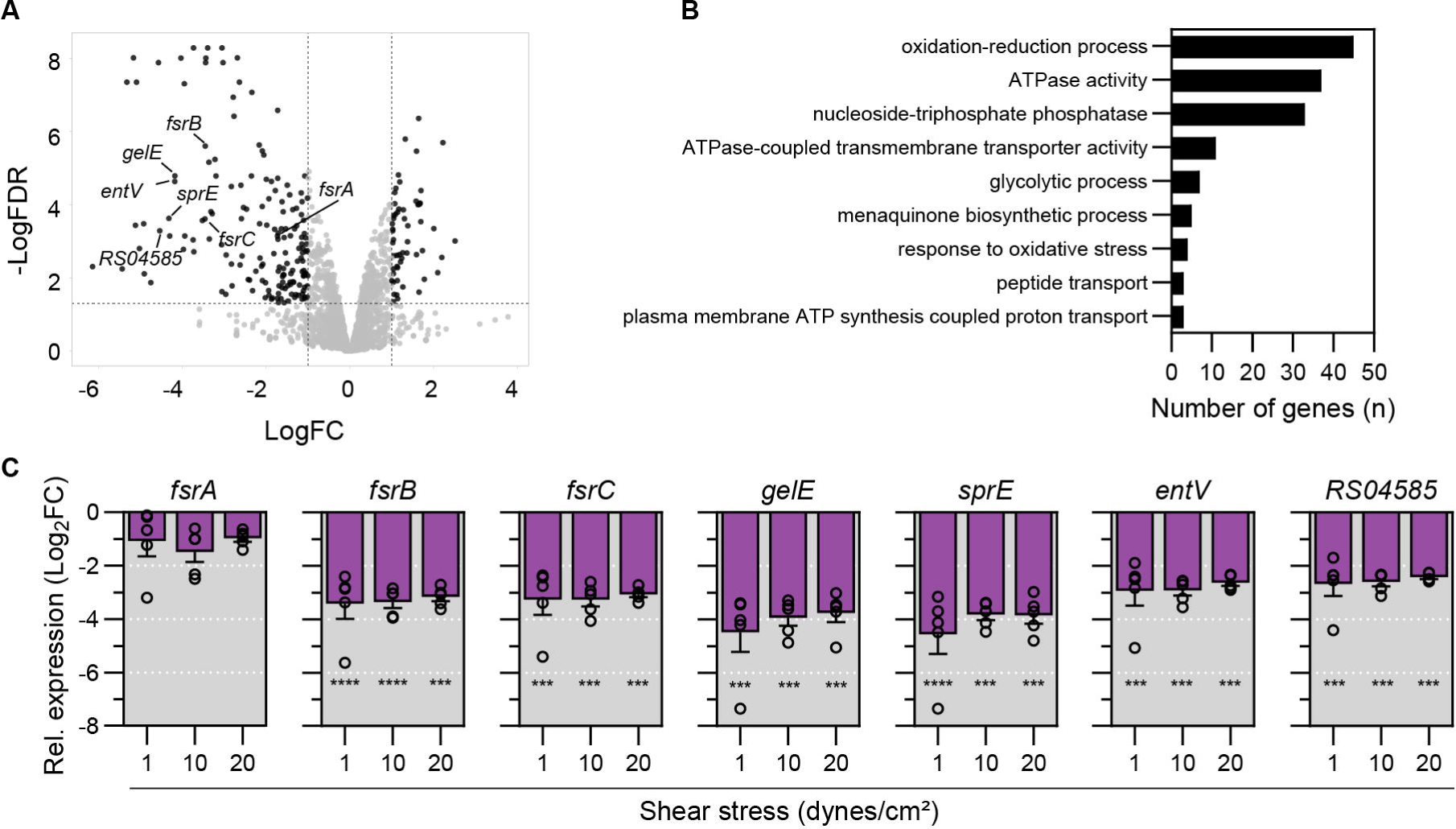
Fluid flow prevents Fsr QS system induction via advection. **A**. Volcano plot showing differentially expressed genes in *E. faecalis* OG1RF after exposure to fluid flow for 30 min compared to static conditions. Genes exhibiting log_2_FC > 1.0 with FDR < 0.05 are shown in black. *fsr* locus genes and associated regulon are labeled. N = 3 independent experiments **B.** Gene Ontology (GO) enrichment analysis showing the most significantly enriched biological processes (p < 0.05) in *E. faecalis* when subjected to fluid flow. **C.** Mean relative gene expression under different magnitudes of shear stress compared to static conditions assessed by qPCR. N = 5, error = SEM. Statistical significance was determined using a one-way ANOVA test and Tukey’s multiple comparison test; *** = p < 0.001, **** = p < 0.0001.

### Early colonization of vegetation is independent of the Fsr QS system

We hypothesized that the combination of advection and low population density at the onset of IE might limit Fsr activation in superficially adhered bacteria. To test this hypothesis, we used a rat model of IE and harvested aortic valve vegetations at 6 hours post-infection (hpi) with either the OG1RF parental wild type or Δ*fsrABDC* strain (hereafter called WT and Δ*fsr* respectively). We found no statistically significant differences in vegetation weight or bacterial colony forming units (CFU) in vegetations or in blood between these two strains (**Fig. 2A-C**). Imaging of vegetations at 6 hpi revealed the presence of single superficial bacteria for both WT and Δ*fsr* strains (**Fig. 2D**). Collectively, these data show that Fsr QS plays a limited role in the superficially and sparsely colonized vegetations of early infection and is dispensable for the establishment of IE. This observation agrees with earlier findings showing no difference in the ability of OG1RF and a *fsrB* deletion mutant to induce IE in a rat model ^48^.

**Fig. 2.**
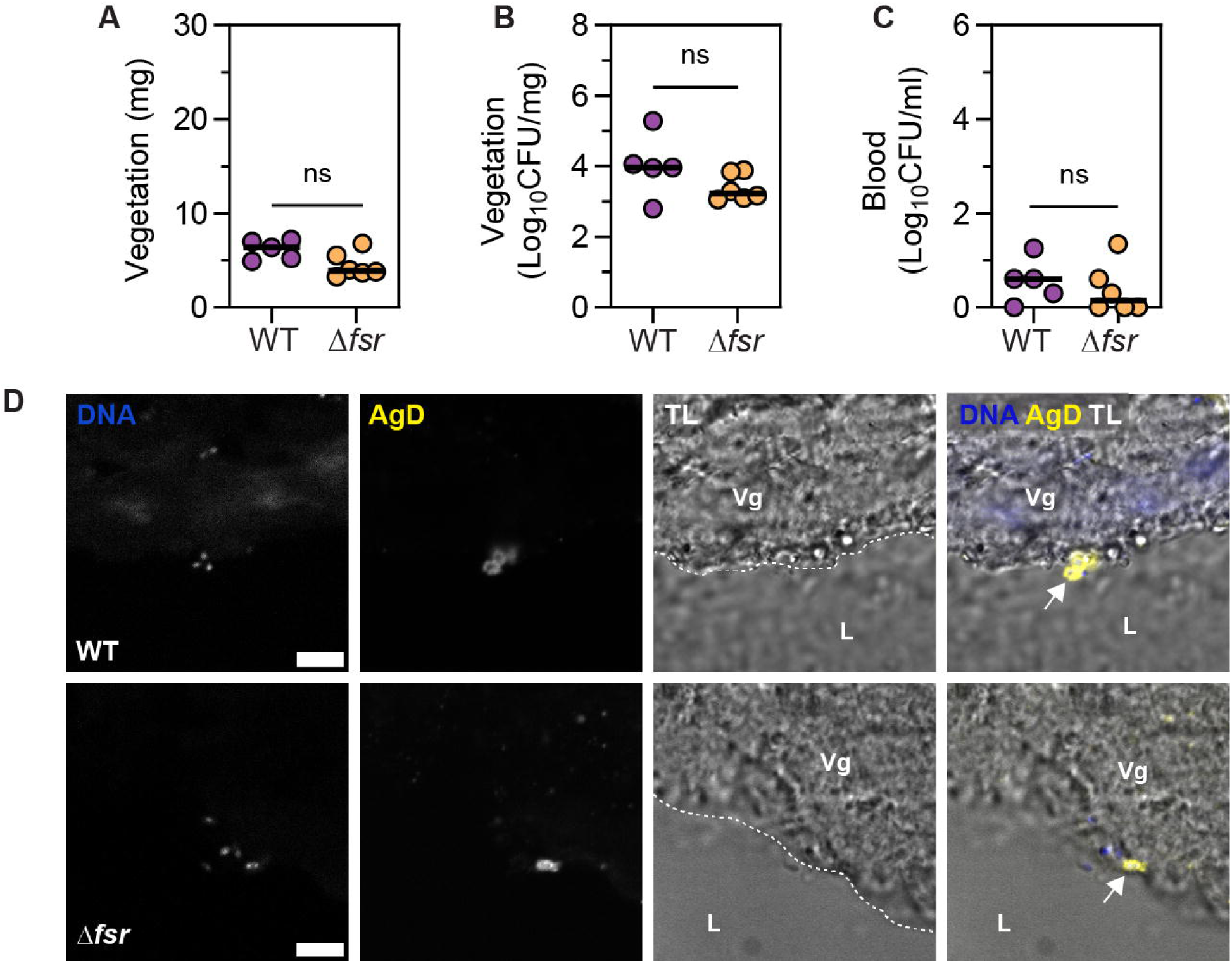
Early colonization of vegetation is independent of the Fsr QS system. **A-C**. Vegetation weight (A), vegetation CFU (B), and blood CFU (C) at 6 hpi. Median of n = 5 - 6 animals per group from N = 1 experiment is shown. Statistical significance was determined using a Mann-Whitney test, ns = not significant. **D**. *E. faecalis* adhesion on vegetation (Vg) surface at 6 hpi captured with LSCM, stained for DNA and *Enterococcus-*specific Group D Streptococcus antigen (AgD) and merged with transmitted light (TL) images. L = lumen, dashed line = vegetation boundary, arrow = bacteria. Representative images are shown from n = 2 animals per group, harvested from N = 2 independent experiments. Scale = 5 µm.

### Absence of Fsr QS system promotes biofilm growth in late IE

As IE progresses, vegetations continue to grow in size through bacterial replication and accumulation of platelets and fibrin. We hypothesized that *E. faecalis* embedded in the interior of mature vegetations might be shielded from blood flow and protected from the effect of advection, thereby favouring Fsr QS activation. We therefore quantified the expression of *fsrC* in WT-infected vegetations harvested from rats at 72 hpi and found that *fsrC* was expressed at higher levels compared to the bacterial inoculum (**Fig. 3A**, p < 0.01), indicating that the intravegetational milieu promotes the activation of QS. To investigate whether *fsr* expression affects virulence at later stages of infection, using the same IE model, we infected rats with the WT and Δ*fsr* strains. We found that 72h vegetations infected with Δ*fsr* were larger compared to WT (**Fig. 3B**, p < 0.05). Moreover, Δ*fsr*-infected animals exhibited a higher bacterial load within each vegetation, compared to WT-infected animals (**Fig. 3C**, p < 0.001)). Immunofluorescence on vegetation sections corroborated this finding (**Fig. 3F**), showing that Δ*fsr* biofilms covered a larger vegetation area compared to WT (**Fig. 3D**, p < 0.01), ranging between 21.87 – 42.10 % for Δ*fsr* compared to 5.19 – 15.35 % for WT. Furthermore, individual Δ*fsr* microcolonies within vegetations exhibited a larger cross-sectional area compared to WT (**Fig. 3E & G**, p < 0.0001). Growth curves comparing WT and Δ*fsr* strains showed no growth advantage for the mutant strain in serum-supplemented BHI (**Fig. S1A**). Gelatinase has been shown to degrade polymerized fibrin ^49^, a main component of vegetations. This degradation has been proposed as a mechanism for bacteria to disperse from the vegetations into the bloodstream^50^, which offered a possible explanation for the increased biofilm growth of the Δ*fsr* strain. However, bacterial CFU in blood, liver, and spleen were similar for both strains, indicating that the increased bacterial burden in vegetations was likely not due to defective dispersal and dissemination (**Fig. S1B-D**). Overall, these data demonstrate that Fsr plays a role in limiting biofilm and vegetation size in late IE.

**Fig. 3.**
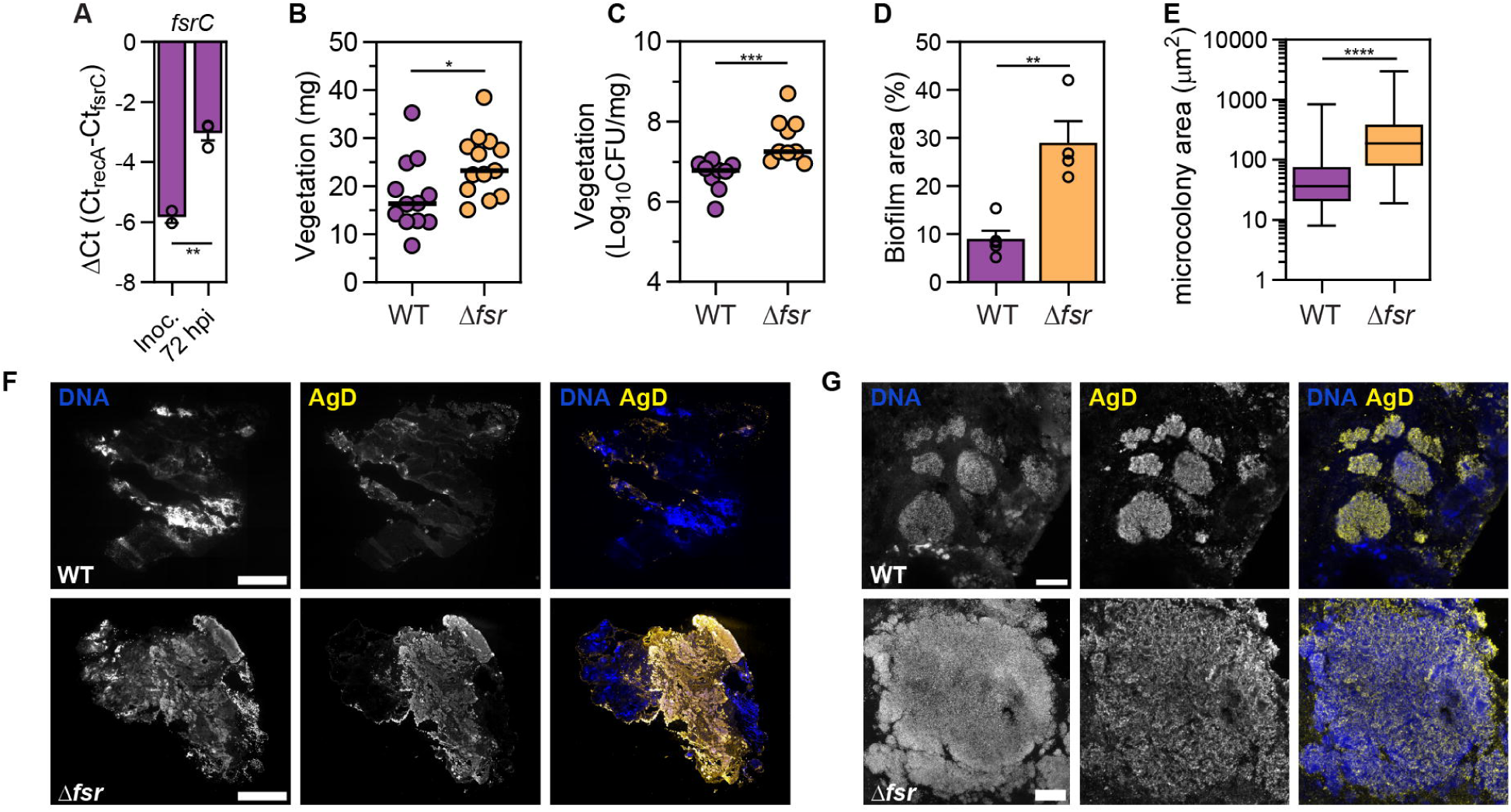
Absence of Fsr QS system promotes biofilm growth in late IE. **A.** qPCR analysis showing mean *fsrC* expression of the WT strain in the inoculum and vegetations at 72 hpi. *fsrC* expression is presented as ΔCt values normalized to the expression of the reference gene *recA*. Vegetation cDNA from n = 3 rats from N = 2 independent experiments was used. Error = SEM. Statistical significance was determined using a t-test. **B.** Vegetation weight at 72 hpi. Median of n = 12-13 animals per group from N = 3 is shown. Statistical significance was determined using a Mann-Whitney test. **C.** Vegetation CFU at 72 hpi. Median of n = 9 animals per group from N = 2 is shown. Statistical significance was determined using a Mann-Whitney test. **D.** Mean biofilm coverage (%) of vegetation sections is shown for n = 4 – 5 animals per group from N = 2. Biofilm coverage was determined by calculating the antigen D (AgD) positive area to total vegetation area. Error = SEM. Statistical significance was determined using a t-test. **E.** Median of **i**ntravegetational microcolony area from n = 2 – 3 animals per group from N = 2 is shown. A total of 88 microcolonies for WT and 101 microcolonies for Δ*fsrABDC* were quantified. Statistical significance was determined using a Mann-Whitney test. **F.** Tile images of whole vegetation sections from 72 hpi captured with fluorescence microscopy, stained for DNA and AgD. Representative sections from n = 4 – 5 animals per group from N = 2 are shown. Scale = 500 µm. **G**. Z-projections of biofilm microcolonies from 72 hpi captured with LSCM stained for DNA and AgD. Representative images from n = 4 – 5 animals per group from N = 2 are shown. Scale = 20 µm; * = p < 0.05, ** = p < 0.01, *** = p < 0.001, **** = p < 0.0001.

### Absence of *gelE* and *sprE* contributes to biofilm growth in late IE

The *fsr* regulon includes *gelE* and *sprE*, which encode the metalloprotease gelatinase and serine protease respectively. These secreted proteases are major virulence factors in *E. faecalis*, affecting cell division, autolysis, adhesion, opsonization, and extracellular matrix composition ^45,49,51–53^. Thus, we hypothesized that Fsr might mediate vegetation and bacterial growth control via gelatinase and/or serine protease activity. As expected for a Fsr-regulated gene, similar to *fsrC* expression (**Fig. 3A**), *gelE* and *sprE* expression was higher at 72 hpi in WT-infected vegetations compared to the inoculum (**Fig. 4A, B**, p < 0.05). When we infected rats with a Δ*gelE* mutant, we found no difference in vegetation weight compared to WT infection (**Fig. 4C**). However, we observed an increase in vegetation CFU for Δ*gelE* (**Fig. 4D**, p < 0.05, Median_Δ*gelE*_ – Median_WT_ = 0.290), albeit less pronounced compared to Δ*fsr* (Median_Δ*fsr*_ – Median_WT_ = 0.468). Growth curves comparing WT and Δ*gelE* strains showed no growth advantage for the mutant strain *in vitro* (**Fig. S1E**). Despite a previously suggested role of gelatinase in promoting biofilm dispersion ^49^, we observed no differences in systemic spread of bacteria between WT and Δ*gelE* (**Fig. S1F-H**). To investigate the spatial organization of the increased bacterial load in Δ*gelE* vegetations, we localized *E. faecalis* within vegetation sections at 72 hpi. Immunofluorescence showed increased biofilm coverage in Δ*gelE*-infected vegetation sections (**Fig. 4E**), with larger individual microcolonies (**Fig. 4F, H**) compared to WT (**Fig. 3E)**. Gelatinase mutant biofilm coverage ranged between 11.49 -38.60 % of the total vegetation area (**Fig. 4G**), whereas WT biofilm coverage ranged between 6.13 – 12.1 %, a similar range to what was observed in independent experiments in comparison with Δ*fsr*- infected vegetations (**Fig. 3D**). Similar to our observation for Δ*gelE* infection, Δ*sprE*-infected vegetations exhibited a comparable weight to WT (**Fig. 4I**) but an increase in CFU (**Fig. 4J**, p < 0.05, Median_Δ*sprE*_ – Median_WT_ = 0.324). Growth curves comparing WT and Δ*sprE* strains showed no growth advantage for the mutant strain during exponential phase *in vitro* (**Fig. S1I**). We observed an increase in spleen CFU in Δ*sprE*-infected animals but no difference in blood and liver CFU compared to WT (**Fig. S1J-L**). Infection with the double deletion mutant Δ*gelE*Δ*sprE* exhibited a similar vegetation weight (**Fig. 4K**) and an increased vegetation CFU compared to WT **(Fig. 4L**, p < 0.05, Median_Δ*gelE*Δ*sprE*_ – Median_WT_ = 0.376). Despite the lack of both proteases, the increase was not as pronounced as in the absence of *fsr*. Growth curve comparison between WT and Δ*gelE*Δ*sprE* showed no difference in growth during exponential phase (**Fig. S1M**). Blood, spleen, and liver CFU were lower compared to WT (**Fig. S1N-P**, p < 0.05), indicating a defect in systemic spread. However, this dispersion defect was absent in Δ*fsr* infection (**Fig. S1B-D)**, suggesting that other factors regulated by Fsr might compensate for impaired dispersal when both *gelE* and *sprE* are downregulated. Collectively, our data show that gelatinase and serine protease contribute partially to biofilm growth control conferred by the *fsr* locus in IE vegetations and suggest that additional *fsr* associated factors likely also contribute.

**Fig 4.**
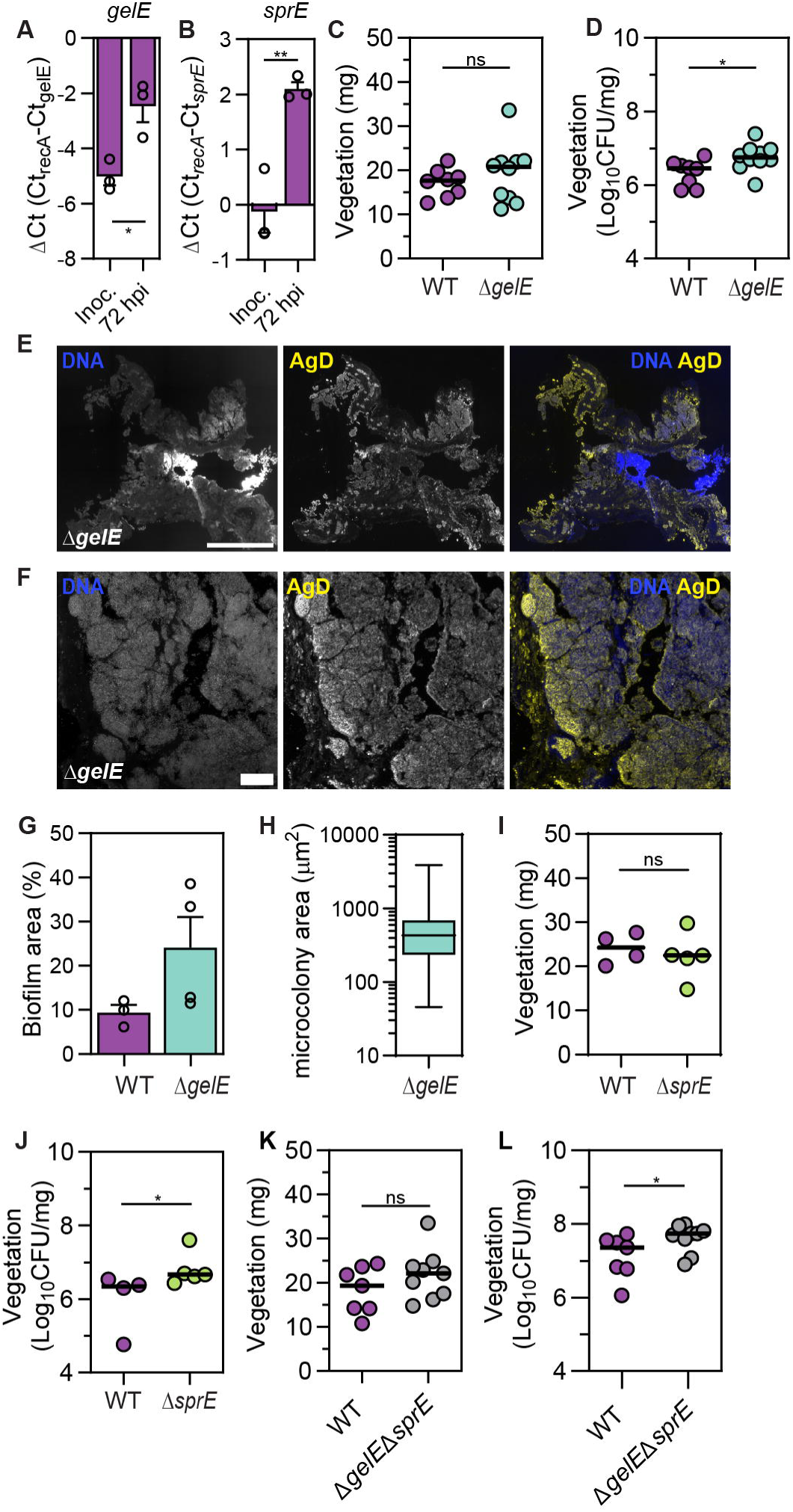
Absence of *gelE* and *sprE* contributes to biofilm growth in late IE. **A-B.** qPCR analysis showing mean *gelE* and *sprE* expression of the WT strain in the inoculum and vegetations at 72 hpi. *gelE* expression is presented as ΔCt values normalized to the expression of *recA*. Vegetation cDNA from n = 3 rats from N = 2 independent experiments was used. Error = SEM, statistical significance was determined using a sample t-test. **C-D.** Vegetation weight (B) and CFU (C) at 72 hpi. Median of n = 8 - 10 animals per group from N = 2 is shown, with statistical significance assessed with a Mann-Whitney test. **E.** Epifluorescence microscopy tile images of whole vegetation sections from 72 hpi, stained for DNA and antigen D (AgD). Representative section from n = 4 animals from N = 2 is shown. Scale = 500 µm. **F**. Z-projection of biofilm microcolonies from 72 hpi captured with LSCM stained for DNA and AgD. Representative image from n = 4 animals from N = 2 is shown. Scale = 20 µm. **G.** Mean biofilm coverage (%) per vegetation section from n = 4 animals from N = 2 is shown. Biofilm coverage was determined by calculating the antigen D positive area to total vegetation area. Error = SEM. **H.** Microcolony area median of n = 1 animal from N = 1. A total of 65 microcolonies were quantified. **I-L**. Vegetation weight and CFU at 72 hpi. Median of n = 4 – 5 animals per group from N = 1 for (H) and (I) and n = 7 – 9 from N = 2 for (J) and (K) is shown, with statistical significance assessed with a Mann-Whitney test; * = p < 0.05, ** = p < 0.01, ns = not significant.

### *E. faecalis* gelatinase activates IL-1β

Dispersal impairment could not explain the increased biofilm coverage observed in the absence of the *fsr* locus, as there were no differences in the systemic spread of bacteria compared to WT infection. Therefore, we hypothesized that the Fsr-negative biofilm might interfere with the immune response, allowing the biofilm to proliferate unchecked. In support of this hypothesis, we observed that the increased bacterial load (measured by *E. faecalis* RpoB) in Δ*fsr*- and Δ*gelE*-infected vegetations was not followed by an increased infiltration of host cells (measured by rat histone H3 and β-actin) compared to WT*-*infected vegetations at 72 hpi (**Fig. S2A-F**). Moreover, absolute quantification of immune cells showed no significant difference for either total leukocytes or the neutrophil subpopulation in *ΔgelE*-infected vegetations compared to WT (**Fig. S2G-I**), suggesting that increased bacterial load does not correspond with increased immune cell infiltration, and that Δ*fsr* and *ΔgelE* biofilms may be less immunostimulatory compared to WT.

We next investigated whether gelatinase triggers inflammation by activating IL-1β, as shown previously for *Streptococcus pyogenes* SpeB and *Pseudomonas aeruginosa* LasB proteases ^54,55^. We detected IL-1β in both WT and Δ*gelE*-infected vegetations at 72 hpi using immunostaining (**Fig. 5A, Fig. S6A**). IL-1β fluorescence was notably stronger at the interface between the biofilm and neutrophils undergoing NETosis (**Fig. 5A, Fig. S3A**), and within incoming neutrophils (**Fig. S3B**). Western blots using the same antibody confirmed the predominance of the IL-1β pro-form (pro- IL-1β) in WT and Δ*fsr* vegetations at 72 hpi (**Fig. 5B**). Given the extracellular abundance of IL-1β in direct contact with IE biofilms, we investigated whether *E. faecalis* gelatinase is involved in the activation of IL-1β. Incubation of human pro-IL-1β with WT, Δ*fsr*, and Δ*gelE* strains revealed gelatinase-dependent cleavage of pro-IL-1β into ∼17 kDa fragments, resembling mature IL-1β following cleavage and activation by caspase-1 ^56^ (**Fig. 5C, Fig. S3B**). A *sprE*-defective strain did not abolish cleaving of pro-IL-1β. Following incubation of pro-IL-1β for 6 h with WT or Δ*gelE*::*gelE*^E352A^, which expresses proteolytically inactive gelatinase, we exposed the cell-free supernatants to an IL-1R reporter cell line that specifically recognizes cleaved and active IL-1β. Only the WT supernatant activated the reporter cells, confirming bioactive IL-1β production (**Fig. 5D**, **Fig. S3C**). Mass spectrometry of IL-1β fragments generated by the WT strain revealed a unique cleavage site between F104 and F105, consistent with known gelatinase substrate specificity ^57^ and analogous to caspase-1 cleavage (**Fig. 5E, F**). These findings demonstrate that *E. faecalis* gelatinase cleaves pro-IL-1β into an active form with the potential to promote host cell infiltration to restrict biofilm growth in IE, providing a plausible explanation for the unchecked biofilm growth in the absence of *fsr* or gelatinase.

**Fig. 5.**
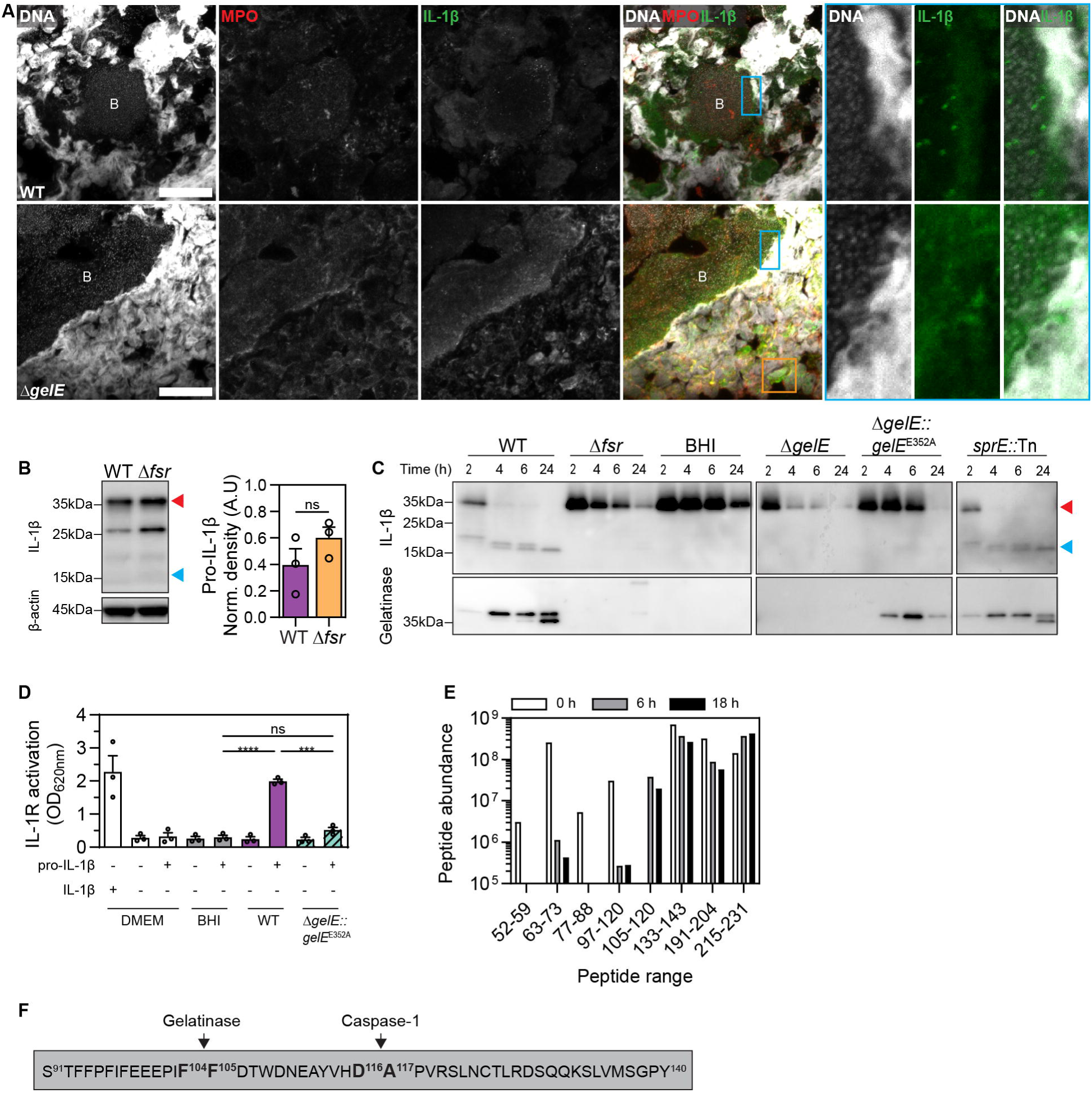
*E. faecalis* gelatinase cleaves and activates IL-1β. **A**. Z-projections of WT and Δ*gelE* vegetations at 72 hpi, captured with LSCM and stained for DNA, myeloperoxidase (MPO), and IL-1β. Cyan insets are highlighting the presence of IL-1β between the NETs-biofilm interface. Orange inset is shown magnified in Fig. S3B, and it is highlighting the colocalization of IL-1β in neutrophils. Representative images from n = 3 animals per group from N = 1 experiment are shown. B = biofilm, scale = 20 µm. **B.** Detection and quantification of IL-1β in WT and Δ*fsr* vegetations at 72 hpi with western blotting. β-actin was used as a loading control. Red arrowhead = pro-IL-1β, blue arrowhead = mature IL-1β. Mean is shown from n = 3 of N = 1. Error = SEM, ns = not significant. Statistical significance was assessed with a t-test. **C.** Pro-IL- 1β and its cleaved fragments in supernatants from OG1RF WT and mutant strains detected by western blotting. Pro-IL-1β was incubated in BHI with indicated OG1RF strains and sampled at 2, 4, 6, and 24 h. Gelatinase presence was also determined by western blotting in these supernatants. Δ*gelE*::*gelE*^E352A^ expresses proteolytically inactive gelatinase. BHI = Negative control with only media and pro-IL 1β. Red arrowhead = pro-IL-1β, blue arrowhead = mature IL-1β. **D.** Activation of HEK-Blue IL-1R reporter cells in the presence of supernatants harvested from OG1RF WT and Δ*gelE*::*gelE*^E352A^ cultures with or without pro-IL-1β at 18 h. Cell activation was assessed by spectrophotometric measurement of reporter cell supernatants incubated with a chromogenic substrate. Stimulation of cells with mature human IL-1β was used as a positive control. N = 3, Error = SEM. Statistical significance was determined with one-way ANOVA. **E.** Mass spectrometry analysis of pro-IL-1β peptide abundance after incubation with OG1RF WT for 0, 6, and 18 h. **F**. Schematic representation of gelatinase and caspase-1 cleavage sites; *** = p < 0.001, **** = p < 0.0001, ns = not significant.

### *fsr* locus absence promotes sugar phosphotransferase system and antiholin-like protein upregulation

Since enhanced biofilm growth in the absence of the *fsr* locus was only partly explained by the absence of gelatinase and serine protease, we hypothesized that the absence of *fsr* might also enhance biofilm growth by optimizing its ability to exploit available nutrients and adapt to the microenvironment of the vegetation. We therefore compared the *E. faecalis* transcriptome of 72 hpi vegetations infected with WT and Δ*fsr* strains. We found 292 genes upregulated and 83 genes downregulated (Log_2_FC ≥ 1, FDR < 0.05) in Δ*fsr* compared to the WT strain (**Fig. 6A, Supplementary File 2**). As expected, the Fsr regulon comprising *gelE* (Log_2_FC = -8.2), *sprE* (Log_2_FC = -8.1), *entV* (Log_2_FC = -7.7), and *OG1RF_RS04585* (Log_2_FC = -8.8) exhibited the most pronounced downregulation. Among the upregulated genes, antiholin-like murein hydrolase modulator *lrgA* (OG1RF_RS12570) and antiholin-like protein *lrgB* (OG1RF_RS12565) exhibited the greatest increase in expression (Log_2_FC = 5.0) (**Fig. 6A**). To identify other biological processes that may be significantly enriched in the absence of the *fsr* locus *in vivo*, we conducted a gene ontology (GO) enrichment analysis. We found that differentially expressed genes were significantly enriched in processes related to nucleotide biosynthesis, sugar transport and carbohydrate metabolism (**Fig. 6B**). One of the most enriched categories was phosphoenolpyruvate-dependent sugar phosphotransferase systems (PTS) systems, which are involved in the uptake and phosphorylation of specific carbohydrates from the extracellular environment. Thus, we concluded that the Fsr QS system likely represses multiple metabolic pathways in vegetations, and its absence leads to the derepression and enhanced utilization of various substrates available in the microenvironment of the vegetation.

**Fig. 6.**
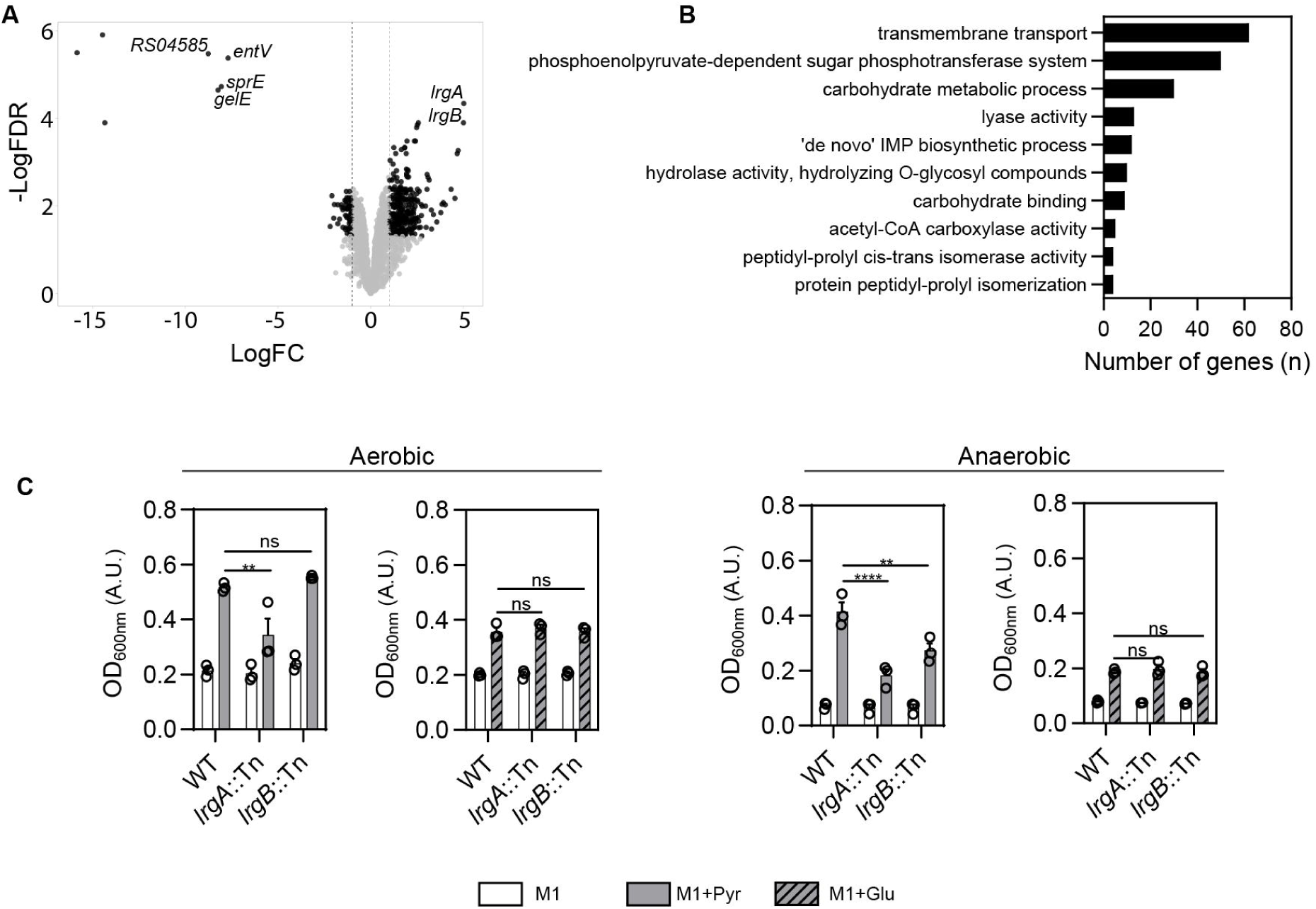
*fsr* locus absence promotes sugar phosphotransferase system and antiholin-like protein upregulation. **A.** Volcano plot showing differentially expressed genes of *E. faecalis* in Δ*fsr*-infected vegetations compared to WT at 72hpi. Genes exhibiting Log_2_FC ≥ 1.0 with FDR < 0.05 are shown in black. *fsr*-associated regulon and *lrgAB* are labeled. n = 4 animals per group from N = 1 experiment. **B.** Gene Ontology (GO) enrichment analysis showing the most significantly enriched biological processes in *E. faecalis* in Δ*fsr*-infected vegetations at 72 hpi. **C.** Endpoint absorbance of OG1RF WT, *lrgA*::Tn, and *lrgB*::Tn grown in M1 media, with or without pyruvate (110mM) or glucose (110mM) supplementation, under aerobic and anaerobic conditions. Mean and SEM is shown for N = 3, two-way ANOVA was applied; ** = p < 0.01, **** = p < 0.0001, ns = not significant

*lrgAB* homologues in *Bacillus subtilis*, *S. aureus*, and *Streptococcus mutans* are involved in transporting pyruvate ^58–61^, which is present in blood ^62^. To investigate whether *lrgAB* plays a similar role in *E. faecalis*, we cultured OG1RF WT, Δ*fsr, lrgA*::Tn, and *lrgB*::Tn strains in minimal medium M1, with and without pyruvate, under both aerobic and anaerobic conditions, to account for potentially different oxygen availability within the large 72-h vegetations. Despite pyruvate supplementation, growth of *lrgA*::Tn was impaired compared to WT under both conditions, with the impairment being more pronounced under anaerobic conditions, whereas the *lrgB*::Tn strain exhibited impairment only under anaerobic conditions (**Fig. 6C**).

These growth defects were specific to pyruvate supplementation, as *lrgA*::Tn and *lrgB*::Tn reached the same growth levels to WT with glucose supplementation (**Fig. 6C**). These findings suggest that *lrgAB* in *E. faecalis* may also be involved in pyruvate transport and utilization. *lrgAB* can restrict phage-mediated extracellular lysis in *E. faecalis* and can inhibit murein hydrolase activity in *S. aureus*, limiting autolysis and conferring tolerance to penicillins ^63,64^. Our preliminary data, however, did not reveal any differences in growth between WT, *lrgA*::Tn, and *lrgB*::Tn when exposed to triton X-100 or ampicillin (**Fig. S4A, Table S1**). We also did not observe any differences in biofilm formation *in vitro* between WT, *lrgA*::Tn, and *lrgB*::Tn (**Fig. S4B, C**). Overall, these data suggests that Fsr QS plays an important role in controlling substrate utilization in vegetations, providing an additional explanation for the increased biofilm formation observed *in vivo*.

### Absence of Fsr correlates with increased antibiotic tolerance, longer bacteraemia, and cases of high disease severity in IE

We hypothesized that the larger biofilm phenotype in Δ*fsr* might contribute to increased tolerance to antibiotic treatment. To test this hypothesis, we treated WT- and Δ*fsr-*infected rats with gentamicin at 48 hpi and harvested the vegetations at 72 hpi. Vegetations presented decreased weight and CFU in the antibiotic-treated WT-infected group (**Fig. 7A, B**, p < 0.05). This decrease correlated with an increase in systemic platelet counts (**Fig. 7C**, p < 0.05), suggesting a reduction in disease severity ^65–67^. By contrast, antibiotic treatment had no effect on vegetation weight and CFU, or platelet counts in Δ*fsr*-infected animals. CFU in blood remained unchanged for both strains, irrespective of antibiotic treatment **(Fig. 7D)**. Microbroth dilution assay showed no difference in gentamicin MIC between the WT and Δ*fsr* strains (**Table S2**). Overall, these data demonstrated that the absence of QS results in a more recalcitrant infection *in vivo*, underscoring a role for Fsr in the antibiotic tolerance of *E. faecalis* biofilms in IE.

**Fig. 7.**
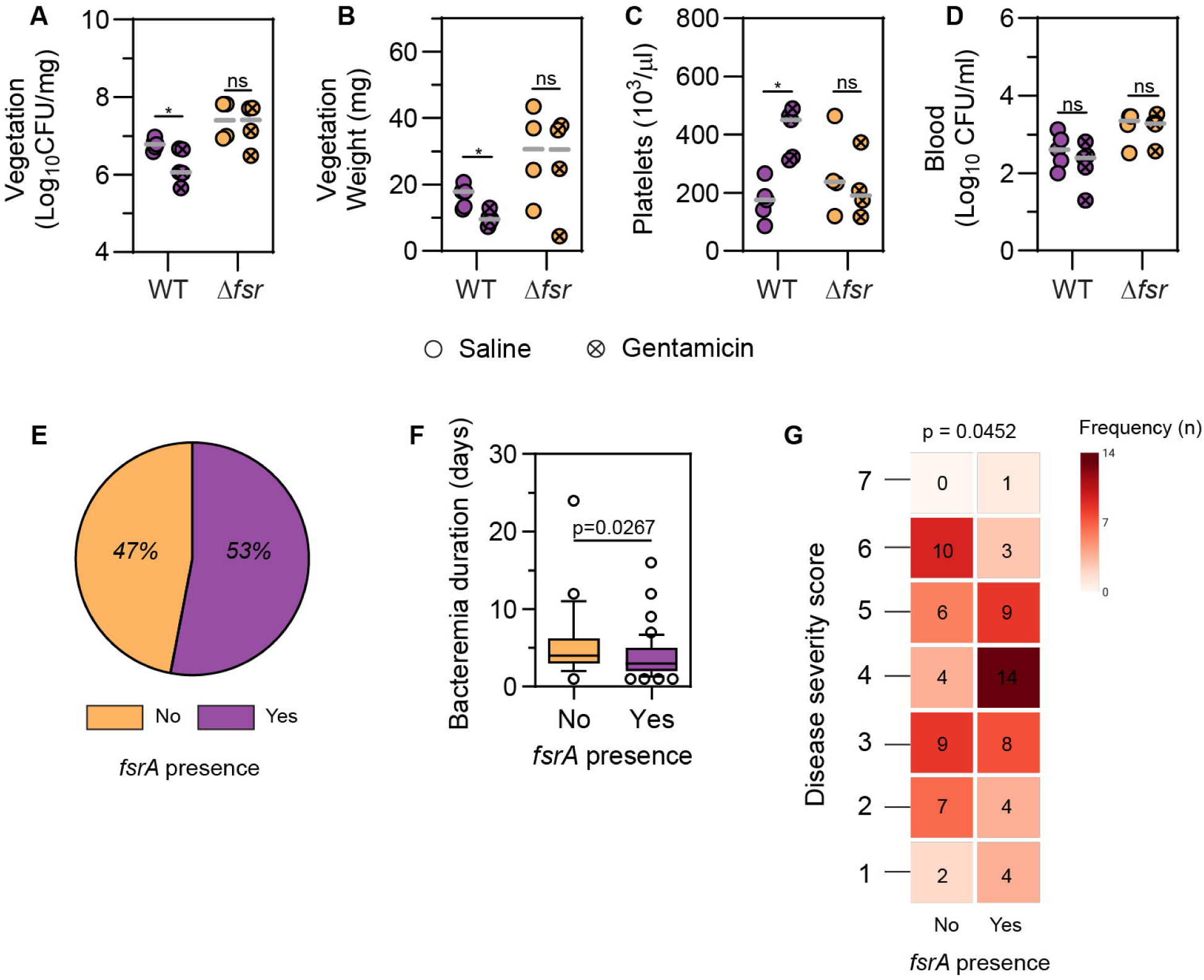
Absence of Fsr correlates with increased antibiotic tolerance, longer bacteraemia, and cases of high disease severity in IE. **A-D.** Vegetation weight (A), vegetation CFU (B), platelet blood concentration (C), blood CFU (D) at 72 hpi, with and without gentamicin treatment. Median of n = 4 – 5 animals per group from N = 1 independent experiment is shown. Statistical significance was determined with a Mann-Whitney test between treated and untreated groups infected with the same strain; * = p < 0.05, ns = not significant. E. *fsrA* presence (%) in *E. faecalis* isolates from IE patients. **F.** Bacteremia duration of IE patients in relation to *fsrA* presence. Median, 10^th^, and 90^th^ percentiles are shown. Statistical significance was determined with Wilcoxon rank sum test with continuity correction. **G.** Cumulative disease severity score (Y-axis) of IE patients infected in relation to *fsrA* presence (X-axis). Frequency (n) for each disease score is shown within each heatmap box. Statistical significance was determined with Fisher’s exact test.

*E. faecalis* clinical isolates often exhibit a chromosomal deletion encompassing the *fsr* locus, resulting in the lack of gelatinase activity ^12,13,68^. In a prior study of 80 IE isolates, 38.7 % showed an absence of gelatinase activity, with 30 % of these lacking the *fsrB* gene ^13^. Although the presence of the *fsr* locus was not associated with disease causation ^13^, its impact on disease severity was not studied. Based on our pre-clinical findings linking *fsr* absence to increased virulence, we hypothesized that there might be a correlation between *fsr* absence and IE severity. To address this, we examined 81 enterococcal IE cases and isolates from cohorts in the United States ^69^ and Switzerland. We first determined the prevalence of *fsrA* as a surrogate for the *fsr* locus. Among the isolates, 47 % lacked *fsrA* (**Fig. 7E**). Clinical data revealed that the absence of *fsrA* was associated with a longer duration of bacteraemia (**Fig. 7F**, p = 0.0267). To better assess disease severity, we implemented an explorative, not clinically validated, scoring system with the objective of obtaining a multidimensional representation that would facilitate a comprehensive understanding of disease severity. The scoring system parameters included fulfilment of the modified Duke criteria for IE ^70^, intensive care unit (ICU) length of stay, need for device or valve replacement, and in-hospital mortality. The correlation of *fsrA* presence with these parameters was examined both individually and within the framework of the scoring system. Although *fsrA* presence was not directly associated with any individual parameter (**Fig. S5**, **Supplementary File 3**), we identified a significant association between high disease severity scores and the absence of *fsrA* (**Fig. 7G**, p = 0.0452). Differences in population structures and treatment regimens between the cohorts were noted (**Supplementary File 3**), but a generalized linear model used to investigate a potential cohort bias in the bacteraemia duration of *fsrA*-positive patients showed no cohort bias (p = 0.104). Overall, these findings suggest that the absence of *fsrA* in *E. faecalis* may correlate to more severe manifestations of IE in human patients.

## DISCUSSION

In this study, we explored the impact of fluid flow on *E. faecalis* physiology under conditions mimicking the heart valve microenvironment. We identified the Fsr QS system as an important negative regulator of biofilm formation in IE. We show that deletion of *fsr* enhances biofilm growth, partly due to *gelE* and *sprE* downregulation. We also demonstrate that gelatinase activates IL-1β, suggesting that its downregulation in *fsr*-negative biofilms may reduce localized inflammation, enabling bacteria to grow unchecked by immune cells. Furthermore, the absence of *fsr* correlates with increased antibiotic tolerance *in vivo*. Building on these preclinical findings, we found that the absence of *fsr* in *E. faecalis* IE isolates correlates with longer bacteremia and high disease severity, providing a possible explanation for the frequent loss of *fsr* among clinical *E. faecalis* isolates.

The Fsr QS system has been previously shown to promote biofilm formation *in vitro* in OG1RF and V583 strains ^53,71,72^ and contribute to disease pathogenesis in endophthalmitis and peritonitis models ^22,73^. Rather than promoting biofilm formation, however, we show that induction of the Fsr QS system *in vivo* limits biofilm growth and reduces pathogenicity in rats. Moreover, we observed that the absence of *fsr* is associated with prolonged bacteremia in patients with IE. Consistent with our findings, the homologous Agr QS system in *S. aureus* also limits biofilm growth and promotes dispersal when activated compared to its inactive state ^74–78^. These contrasting observations in *E. faecalis* likely arise from the dynamic and heterogeneous vegetation microenvironment, which is markedly different from the *in vitro* static conditions commonly used to assess biofilm formation. Besides the heart valve niche, fluid flow is expected to significantly impact QS in other hydrodynamically challenged host niches, including the bladder during intermittent urine expulsion and slow peristalsis in the gut during colonization and disease. At the same time, these niches exhibit complex topographies that could shield biofilm from flow and facilitate localized accumulation of autoinducers. Such spatial effects, previously shown for *S. aureus* and *V. cholerae* biofilms *in vitro* using microfluidics ^74^, could activate QS only in specific regions. Given the direct exposure to blood flow on the endocardial vegetation surface, we propose that adopting a QS-OFF state might be advantageous for *E. faecalis* to rapidly grow into a biofilm and withstand shear stress.

The intact *fsr* locus is variably present among *E. faecalis* strains isolated from IE, blood, urine, and feces ^12,13^. Despite its variable prevalence, its role in disease severity has not been previously explored. Here, we show that the absence of the *fsr* locus correlates with larger biofilms and greater antibiotic tolerance *in vivo* and is also associated with prolonged bacteremia. *S. aureus* Agr-negative isolates have also been associated with persistent bacteremia and higher mortality ^79,80^, often displaying a fitness advantage in the presence of antibiotics *in vitro* ^80^. What drives loss of *fsr* in *E. faecalis* and whether *fsr-*negative strains initiate IE infection or are the product of an in-host adaptation remains unclear. Unlike frameshift mutations in the *agr* locus of *S. aureus* ^81^, loss of *fsr* is typically associated with the deletion of the *fsrC*-EF_1841 region ^68^. *E. faecalis* strains lacking this region harbor a highly conserved ∼600bp junction sequence, suggesting that the region spanning the deletion might be a product of horizontal gene transfer ^12,68^. Since strain transmission from IE patients is not typically expected, and IE is likely an evolutionary “dead end” for the infecting strain, further investigation is needed to identify the conditions promoting loss of *fsr*. Nevertheless, given the fitness advantage of the *fsr* mutant *in vivo*, it remains possible that in-host adaptation could be driving the emergence of QS-deficient cheaters during IE.

IL-1β is a potent pro-inflammatory cytokine typically cleaved and activated intracellularly by caspase-1 upon induction of pyroptosis ^82^, but can also be activated by neutrophil elastase and cathepsin G ^83,84^. In addition to host activation, IL-1β can also be activated by secreted bacterial proteases such as *Streptococcus pyogenes* SpeB and *Pseudomonas aeruginosa* LasB, and IL-1β cleavage has been suggested to serve as a sensor for microbial proteases ^54,55^. Here, we show that *E. faecalis* gelatinase proteolytically activates IL-1β, which is abundant in proximity to vegetation biofilms and is likely released by neutrophils undergoing NETosis in response to bacterial infection. Although gelatinase may activate IL-1β in this context, host proteases present in vegetations are also likely to contribute to IL-1β activation ^83,84^. Rather than serving as the sole activator, gelatinase activation of IL-1β could complement existing host activation mechanisms, thereby amplifying spatially restricted inflammation around the vegetation.

In addition to its role in regulating biofilm via *gelE* and *sprE*, we show that loss of Fsr QS also leads to a large-scale transcriptional reprogramming in *E. faecalis*, suggestive of a shift in metabolic capacity and a much larger (albeit likely indirect) *fsr* regulon than previously appreciated. Genes *lrgA* encoding a murein hydrolase regulator and *lrgB* encoding an anti- holin were the most significantly upregulated genes within Δ*fsr* vegetations. We found that *lrgA and lrgB* are important for *E. faecalis* to grow when pyruvate was the main carbohydrate source under both aerobic and anaerobic conditions *in vitro*, consistent with previous studies showing that the *lrgAB* homologs in *B. subtilis*, *S. aureus*, and *S. mutans* are involved in pyruvate transport ^58–61^. Loss of the *fsr* locus in *E. faecalis* also resulted in the upregulation of several PTS in IE vegetations. Since *E. faecalis* is non-motile, upregulation of PTS might enhance the utilization of nutrients available in their immediate microenvironment. PTS can also regulate numerous cellular processes, including antibiotic tolerance, through the direct phosphorylation of target proteins or by interacting with them in a phosphorylation- dependent manner ^85,86^. Taken together, these data suggest that QS-regulated expansion of metabolic capacities may contribute to the enhanced growth of *E. faecalis* biofilms observed in IE.

Overall, our findings show that the Fsr QS system both restricts biofilm expansion and activates pro-inflammatory signaling in IE. Consequently, spontaneous deletion of the *fsr* locus, as seen in clinical isolates, or possible downregulation of *fsr* expression during the course of infection, simultaneously promotes unchecked biofilm growth with reduced immune activation and may improve metabolic efficiency to support biofilm expansion within the vegetation. Although QS inhibition is pursued as an anti-virulence strategy for a number of pathogens ^87^, our data suggest that suppressing *fsr* may not be beneficial in IE. Additional studies are warranted to evaluate whether the absence of the *fsr* locus or gelatinase function can serve as a prognostic marker for disease severity, potentially guiding more personalized treatment approaches in patients with IE. Collectively, these findings offer a deeper understanding of the infection-associated biofilm physiology of *E. faecalis* and its role in antibiotic tolerance and immune evasion, which could provide insight into other biofilm-related infections and inform future strategies for their treatment.

## METHODS

### Materials

All reagents and instruments used for RNA isolation, SDS-PAGE, and western blotting were purchased from Thermo Fisher Scientific, unless otherwise specified. Antibodies and respective dilutions used for immunofluorescence (IF) and western blotting (WB) in this study include rabbit anti-Streptococcus Group D (Antigen D) (1:500 for IF, Cat. Nr. 12-6231D, American Research Products, Inc), rabbit anti-histone H3 (citrulline R2 + R8 + R17) (1:500 for IF and 1:1000 for WB, Cat. Nr. ab5103, abcam), rabbit anti-histone H3 (1:2000 for WB, Cat. Nr. ab1791, abcam), mouse anti-myeloperoxidase (1:50 for IF, Cat. Nr. NBP1-51148, Novus Biologicals), rabbit anti-IL-1β (1:50 for IF and 1:1000 for WB, Cat. Nr. ab283818, abcam), rabbit anti-gelE (1:500 for WB, Cat. Nr. PA5-117682, Invitrogen), rabbit anti-beta-actin (1:4000 for WB, Cat. Nr. ab8227, abcam), mouse anti-RpoB (1:5000 for WB, Cat. Nr. 663903, Biolegend), goat anti-Rabbit IgG (H+L) HRP-conjugated antibody (1:5000 for WB, Cat. Nr. 31460, Invitrogen), goat anti-mouse IgG (H+L) HRP-conjugated antibody (1:5000 for WB, Cat. Nr. Ab6789, abcam), goat anti-rabbit IgG Alexa 488 (1:1000 for IF, Cat. Nr. A11034, Invitrogen), goat anti-mouse IgG1 Alexa 633 (1:1000 for IF, Cat. Nr. A-21126, Invitrogen).

### Bacterial strains

Strains used in *in vitro* and *in vivo* assays are summarized in **Table S3**. Clinical data and 24 *E. faecalis* isolates from IE patients were collected in accordance with the ethical guidelines of the independent ethics committee Zurich, Switzerland. The study was approved under BASEC ID 2017-01140 and 2017-02225. Informed consent was obtained from all participants prior to data collection. Previously published clinical data and genomic information from 57 *E. faecalis* isolates collected from patients with definite or probable IE at University of Pittsburgh Medical Center were also included ^69^. This study was approved with a waiver of informed consent by the Institutional Review Board at the University of Pittsburgh (protocol no. STUDY22050046).

### Bacterial cultures

All strains were routinely grown in brain heart infusion (BHI) broth at 37°C without shaking, in ambient air, unless otherwise stated. Growth curves were generated by diluting overnight cultures 1:100 in fresh BHI supplemented with 40 % horse serum (Sigma-Aldrich) (BHIS) and adding 100 µl of the diluted cultures in triplicate to a 96-well plate. The plate was then incubated for 18 h at 37°C in a plate reader (TECAN), with absorbance recorded at 600 nm every 20 min in ambient air conditions without shaking. To investigate the role of *lrgAB*, overnight cultures were resuspended 1:100 in M1 media^88^ supplemented with pyruvate (110 mM) or glucose (110 mM) or resuspended in BHI or BHI with 0.4 % Triton X-100 and incubated in ambient air or anaerobic conditions for 24 h at 37°C without shaking with absorbance recorded at 600 nm every 20 min.

### Flow rate calculation

Shear stress on the aortic valves at a normal resting heart rate ranges from 10 to 28.5 dynes/cm², based on experimental and computational data ^42–44^. To mimic the microenvironment of the aortic valves in our *in vitro* assays, we exposed bacteria to 20 dynes/cm^2^, approximating the mean value of this range. To achieve 20 dynes/cm^2^ in microfluidic channels (µ-Slide VI 0.4, ibidi), we calculated the required flow rate using the following formula provided by the manufacturer:

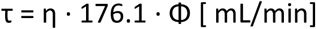

where τ = shear stress (dynes/cm^2^), η = dynamical viscosity (dynes × s/cm^2^) and Φ = flow rate. Dynamic viscosity of BHIS media is 0.0101171 dyn × s/cm^2^ at 37°C, as determined using a viscometer. Thus, the required flow rate to achieve 20 dynes/cm² is 11.226 mL/min.

### Incubation of bacteria under fluid flow conditions

To assess the early impact of fluid flow on *E. faecalis*, 30 µl of stationary-phase bacterial cultures of OG1RF WT were seeded into microfluidic channels with uncoated polymer surface (µ-Slide VI 0.4, ibidi) and incubated for 30 min at 37°C to allow bacterial attachment. After the 30-minute incubation, media in the microchannels was exchanged with fresh BHIS to remove non-attached bacteria, as BHIS approximates *in vivo* growth conditions ^83,84^. For flow conditions, bacteria were exposed to BHIS at defined shear stress levels: low (1 dynes/cm^2^), intermediate (10 dynes/cm^2^), and high shear stress (20 dynes/cm^2^), the latter corresponding to the aortic valve shear stress levels. Pulsatile flow was applied using a peristaltic pump (IPC 12 model, ISMATEC), programmed to mimic cardiac diastole and systole by alternating between pauses of 0.5 s and BHIS delivery for 0.3 s for a total duration of 30 min at 37°C. For static conditions, bacteria were incubated inside the microchannels for 30 min at 37°C without fluid flow. After the incubation period, BHIS in the microchannels was replaced immediately with RNAprotect Bacteria reagent (Qiagen), followed by incubation for 5 min at room temperature (RT). Bacteria were then detached from the microchannels using a 1 mL syringe prefilled with 200 µL of RNAprotect by applying rapid plunging movements to shear bacteria from the surface. Detached bacteria were pelleted at 8,000 g for 15 min at RT and stored in - 80°C until RNA isolation.

### RNA isolation

Bacterial samples harvested from microfluidic channels and treated with RNAprotect or vegetations stored in RNALater Stabilization Solution were transferred in 1 mL Trizol and homogenized with lysing matrix B (MP Biomedicals) at 6 m/s for 3 cycles of 40 s with ice cooling between cycles. Chloroform was added to Trizol at a 1:5 ratio, mixed by gentle inversion, and incubated for 2 min at RT. Following centrifugation at 12,000 g for 15 min at 4°C, the aqueous phase was mixed with an equal volume of 80% ethanol. Samples were loaded onto a RNeasy Mini spin column (Qiagen), and total RNA was purified according to the manufacturer’s instructions, including an on-column DNase treatment step. RNA quantity was measured using the Qubit RNA Broad Range Assay Kit, and genomic DNA contamination was assessed using the Qubit dsDNA HS Assay Kit. RNA quality was evaluated using the RNA ScreenTape and RNA Sample Buffer on a TapeStation 2200 instrument (Agilent), following the manufacturer’s instructions.

### Transcriptomic sequencing analysis

For bacterial total RNA samples isolated from fluid flow conditions, library preparation was performed using the TruSeq Stranded mRNA Library Prep Kit (Illumina). The libraries from fluid flow experiments were sequenced on an Illumina HiSeq 2500 platform to generate 75 bp paired-end reads. Vegetation total RNA samples were processed using the Ribo-Zero Plus rRNA depletion kit (Illumina, USA) to remove rat and bacterial rRNA, followed by library preparation with the TruSeq Stranded mRNA Library Prep Kit (Illumina). These libraries were sequenced on an Illumina Illumina HiSeqX v2.5 platform to generate 150 bp paired-end reads. All data were analyzed using a pipeline on the Galaxy platform ^89^. Briefly, quality control of the sequencing data was performed using FastQC, followed by filtering with SortMeRNA (Version 4.3.6) to remove rRNA sequences. Reads were aligned to the *E. faecalis* OG1RF reference genome (Accession number: GCF_000172575.2_ASM17257v2) using Bowtie2 (Version 2.5.3) ^90^ and reads per gene were quantified using HTSeq-count (Version 2.0.5). Differential expression analysis was conducted with edgeR (Version 3.36.0) with statistical significance set at FDR < 0.05. GO enrichment analysis was performed using goseq (Version 1.50.0).

### Quantitative PCR

cDNA was synthesized from total RNA using the SuperScript III First-Strand Synthesis SuperMix (Thermo Fisher Scientific) according to the manufacturer’s instructions, including a minus reverse transcriptase control for each sample. qPCR primer efficiency was determined by performing qPCRs on 10-fold serial dilutions of pooled cDNA (1:2 to 1:2000) and calculating efficiency (%) using StepOne software (version 2.3, Applied Biosystems). Primer pairs with efficiency 90-100% were selected for subsequent qPCR experiments. All qPCR reactions were performed in duplicates, including minus reverse transcriptase and no-template controls, using the KAPA SYBR FAST qPCR Master Mix (2X) kit (KAPA BIOSYSTEMS) on a StepOnePlus Real-Time PCR System (Applied Biosystems). Data were analyzed using the 2^-ΔΔCt^ method and presented either as ΔCt values or as log2 fold change (log2FC) relative to the control group. To normalize gene expression in bacterial samples exposed to fluid flow, the geometric mean of *recA* and *dnaB* was used, based on their stable expression under both static and fluid flow conditions as indicated by transcriptomic data and validation using the Bestkeeper v1 analysis tool ^91^. To normalize gene expression in vegetation samples, *recA* was chosen as the reference gene due to its stable expression both *in vivo* and *in vitro*, as demonstrated by prior transcriptomic data. Primers used are summarized in **Table S4**.

### DNA manipulation and construction of deletion mutants

Generation of *E. faecalis* knockout mutants was done by allelic replacement using a temperature-sensitive shuttle vector previously described ^92^. Vector pGCP213 was linearized using appropriate restriction enzymes (New England Biolabs, USA) for the construction of the deletion mutants. The flanking regions were fused together by overlap extension PCR to generate the final insert. Linearized vector and inserts were then ligated using the T4 ligase or In-Fusion HD cloning kit (Clontech, TaKaRa, Japan) and transformed into Stellar competent cells. Successful plasmid constructs were verified by Sanger sequencing and subsequently extracted and transformed into OG1RF. Transformants were selected with erythromycin (25 μg/mL) at 30°C and then passaged at a nonpermissive temperature at 42°C with erythromycin to select for bacteria with successful plasmid integration into the chromosome. For plasmid excision, bacteria were serially passaged at 37°C without erythromycin for erythromycin-sensitive colonies. These colonies were then subjected to PCR screening to identify successful deletion mutants. All primers used are listed in **Table S4.** The Δ*fsr* strain, as well as the previously constructed Δ*gelE* strain ^93^, did not show any additional deletions or major mutations compared to WT based on assessment with whole genome sequencing.

The proteolytically inactive GelE strain was generated by chromosomal insertion of gelE*(E352A), which has an E352A point mutation in the active site ^94^ and an A29 silent mutation to introduce a SacII restriction site as a unique identifier of engineered GelE strains ^95^, into *E. faecalis* OG1RF Δ*gelE* ^96^. The silent mutation was first incorporated by amplifying WT gelE sequence from OG1RF genomic DNA, including ∼0.5 kb upstream and downstream regions and 15bp overhangs homologous to vector ends, using the primer pairs gelE_ins_1/gelE_ins_2 and gelE_ins_3/gelE_ins_4, which were subsequently fused by overlap extension PCR to generate the final insert. pGCP213 was linearised by inverse PCR using pGCP213_F and pGCP213_R. The insert and linearised pGCP213 were then ligated by In- Fusion HD Cloning Kit (Clontech, TaKaRa, Japan) and transformed to *E. coli* Stellar cells as described above, generating pGCP213::*gelE*\*. pGCP213::*gelE*\* was then subjected to site directed mutagenesis (SDM) by PCR using overlapping mutagenic primers gelE_E352A_F/gelE_E352A_R. The PCR product was treated with DpnI (New England Biolabs, USA), purified and re-ligated with In-Fusion HD Cloning Kit, generating pGCP213::*gelE* (E352A). The final plasmid was similarly propagated in *E. coli* Stellar cells and subsequently electroporated into Δ*gelE* OG1RF for serial passaging and chromosomal integration. Secretion of inactive gelatinase was validated by western blot of cell-free culture supernatants and gelatinase assay on Todd-Hewitt agar + 3% gelatine ^45^.

For the plasmid complementation of GelE, promoter-free *gelE* sequence from wildtype OG1RF was amplified with primers gelE_pTCV-Ptet_F/gelE_pTCV-Ptet_R (**Table S4**). The *gelE* insert is ligated to a BamHI/PstI-digested pTCV-P_tet_ shuttle vector using InFusion HD Cloning Kit (ClonTech, TaKaRa, Japan) and transformed into Stellar competent cells, generating pTCV-P_tet_::*gelE*. Verified pTCV-P_tet_::*gelE* constructs were extracted, transformed into *E. faecalis* OG1RF Δ*gelE* via electroporation and selected on BHI agar with 25 μg/mL erythromycin. Correct transformants were subsequently maintained in erythromycin (25 μg/mL) for complementation experiments.

### Infective Endocarditis rat model

Male Sprague-Dawley rats (7-9 weeks old) were used in accordance with the NTU Institutional Animal Care and Use Committee guidelines (Animal utilization protocols A19091 and A24076). Animals were kept in cages enriched with bedding and wooden blocks. Food and water were provided at libitum. Infective endocarditis was established as previously described with minor modifications ^97^. Briefly, polytetrafluoroethylene (PTFE) catheters (0.6 mm outer diameter, 6.5 cm in length), with a silicon ring positioned at 5 cm, were sealed at both ends with 28-gauge stainless steel plugs (Braintree Scientific) and sterilized by autoclaving. Aortic valve lesions were induced by catheterizing the left ventricle via the right carotid artery of isoflurane-anesthetized rats. Proper placement of the catheter was assessed by momentary disturbance of the blood flow pattern monitored by a rat paw pulse oximeter (Physiosuite, Kent Scientific) and by the vigorous pulsation of the catheter once inserted in the left ventricle. Catheter was secured in place by tying a silk suture (size 7-0) around the carotid artery and the rostral end of the silicon ring. IE was induced by injecting a 200-µL bacterial suspension (2 x 10^6^ CFU) in PBS via the dorsal penile vein at 24 h post-catheterization. To test antibiotic efficacy, gentamicin (3mg/kg of body weight) or saline were injected via the dorsal penile vein at 48 hpi. Analgesia was provided daily by administering buprenorphine (0.5 mg/kg) subcutaneously. At 6, 24, or 72 hpi, rats were anesthetized with isoflurane inhalation and sacrificed in a CO_2_ chamber, followed by cervical dislocation. The heart was extracted and catheter placement inside the left ventricle was verified. Animals with incorrect catheter placement were excluded from downstream analysis.

For CFU quantification in tissue samples, valvular vegetations, liver, and spleen were weighed and homogenized in a lysing matrix M tube (Cat. Nr. 116923050-CF, MP Biomedicals) with PBS using a FastPrep24 instrument (3 cycles at 4 m/s for 20s each). Homogenates were serially diluted, plated using the drop plate method on BHI agar with and without rifampicin (25 µg/mL), and incubated for 18 h at 37°C.

Blood samples were collected via heart puncture under isoflurane anaesthesia, followed by euthanasia in CO_2_ chamber and cervical dislocation. Blood was transferred in EDTA-coated tubes to prevent coagulation and analyzed using a XN-10 Automated Hematology Analyzer (SYSMEX) at the Advanced Molecular Pathology Laboratory (AMPL) at the Agency for Science, Technology and Research, Singapore. For CFU quantification, 100-500 µL of blood were plated on BHI agar with and without rifampicin (25 µg/mL).

### Tissue immunofluorescence staining

Aortic valve vegetations for immunofluorescence staining were routinely collected and fixed in 4 % paraformaldehyde for 24 h at 4°C, followed by sequential cryoprotection in 15 and 30 % sucrose in PBS at 4°C until the samples sank. Samples were embedded in optimal cutting temperature (OCT) compound (Tissue-Tek, SAKURA Finetek), snap-frozen in a dry ice isopropanol bath, and sectioned at a thickness of 5-8 µm using a cryostat (CM1950, Leica). Vegetations intended for IL-1β staining were embedded in OCT and snap-frozen immediately after harvesting to prevent antigen masking, then sectioned at a thickness of 7 µm and fixed in 4 % paraformaldehyde for 10 min at 4°C. All sections were permeabilized in 0.5 % Triton X-100 for 5 min and blocked with buffer containing 5 % goat serum, 5 % bovine serum albumin (BSA), and 0.05 % Triton X-100 in PBS for 30 min at RT. Sections were incubated overnight at 4°C with primary antibodies diluted in 1 % goat serum, 1 % BSA, and 0.05 % Triton X-100 in PBS. After washing 3 times with PBS, sections were incubated with fluorophore-conjugated secondary antibodies for 1 h at RT in the dark. Nuclei were counterstained with 4’,6- diamidino-2-phenylindole (DAPI) (1:500, stock concentration 5 mg/mL, Thermo Fisher Scientific) for 5 min at RT. Sections were mounted with antifade mounting medium and stored in -20°C until microscopy imaging. Secondary-only antibody controls for goat anti-rabbit IgG Alexa 488 (Cat. Nr. A11034, Invitrogen) and goat anti-mouse IgG1 Alexa 633 (1:1000, Cat. Nr. A-21126, Invitrogen) are shown in **Fig. S6.** Antibodies used are listed in the Materials subsection. All microscopy images were acquired at NTU Optical Bio-Imaging Centre (NOBIC) imaging facilities at SCELSE.

### Microscopy and image analysis

To obtain images of whole vegetation cross-sections, tile epifluorescent images were captured on a Carl Zeiss Axio Observer Z1 inverted microscope equipped with a 20x objective using ZEN 2 (blue) software (Zeiss). Snapshots or Z-stacks were acquired on a Carl Zeiss LSM 780 laser scanning confocal using ZEN 2.3 SP1 FP3 (black) software (Zeiss). Imaging parameters were kept constant across each experiment. Maximum intensity projections were generated from Z-stacks using ImageJ (version 1.54f). Linear brightness and contrast adjustments were applied for visualization purposes where necessary using ImageJ.

Tile epifluorescent images of whole vegetation cross-sections were thresholded to determine the biofilm and vegetation outlines based on antigen D and DAPI fluorescent signals respectively. Based on these outlines, the total biofilm and vegetation area was measured using ImageJ and biofilm coverage (%) was calculated using the formula: (biofilm area/total vegetation area) x 100. For each independent vegetation sample, biofilm coverage (%) was averaged from 3 cross-sections 80 µm apart, which served as technical replicates. Individual biofilm microcolony area was quantified using ImageJ in snapshots obtained at 63x with LSCM, across 3 cross-sections spaced 80 µm apart, from 1-3 vegetations per condition. A total of 50-100 microcolonies were quantified per vegetation.

### Western Blotting

Aortic valve vegetations were homogenized using a combination of lysing matrix M and B (MP Biomedicals) to ensure lysis of both rat and bacterial cells. Vegetations were placed in tubes with the lysing matrices and filled with RIPA lysis buffer (250 μL/5 mg tissue), supplemented with 3x Halt Protease Inhibitor Cocktail and 1 x EDTA. Homogenization was performed using a FastPrep24 instrument (MP Biomedicals) for 4 rounds at 4 m/s, 20 s each. Samples were then incubated on ice for 30 min and sonicated for 3 rounds (15 s on, 5 s off) on ice at 20 % power using a sonicator (Vibra cell, SONICS). Lysates were centrifuged at 10,000 x g for 20 min at 4°C, after which the supernatant with the soluble proteins was collected. Total protein concentration was measured using the Pierce BCA protein assay kit. 25 or 50 µg of vegetation protein for detecting rat only or rat and bacterial targets respectively, and 10 µL of supernatant from IL-1β cleaving assays were separated by SDS-PAGE on NuPAGE 4-12% Bis-Tris gels in ice-cold 1x MES or 1 x MOPS SDS Running Buffer. Electrophoresis was performed at 120 V for 40-60 min. Proteins were transferred to polyvinylidene fluoride (PVDF) membranes using the iBlot®2 Gel Transfer Device. Membranes were blocked with 5 % BSA in 1 X Tris-Buffered Saline with 0.1 % Tween 20 (TBST) for 1 h at RT with shaking. Membranes were incubated with primary antibodies diluted in 1 % BSA in TBST overnight at 4°C with gentle shaking. Membranes were washed 3 times in TBST and incubated with HRP-conjugated secondary antibody diluted in 1 % BSA in TBST for 1 h at RT. Protein bands were visualized using SuperSignal West Femto Maximum Sensitivity Substrate and imaged in Amersham ImageQuant 800. For reprobing, membrane-bound primary and secondary antibodies were removed using Restore PLUS Western Blot Stripping Buffer by incubating the membrane for 30 min at 50°C with shaking. Protein band density was quantified using ImageJ. Antibodies used are listed in the Materials subsection.

### Flow cytometry

Aortic valve vegetations were excised, weighed, and minced in 500 µL of digestion buffer (0.1 mg/mL Liberase in PBS). Samples were incubated at 37°C for 30 min with gentle rocking. After incubation, 500 µL of flow cytometry staining buffer (FCSB; 1 % BSA and 0.1% sodium azide in PBS) was added, and the supernatants were filtered through a 35-µm cell strainer into a 50 mL tube. An additional 500 µL of FCSB was used to flush the strainer, bringing the total volume to 1.5 mL. Filtrates were centrifuged at 500 × g for 5 min at 4°C, and cell pellets were resuspended in 200 µL of FCSB. 100 µL per sample were subsequently used for each staining condition. Non-specific Fc receptor binding was blocked by incubating with mouse anti-CD32 (1:100, Cat. Nr. 550270, BD Pharmingen) for 20 min at 4°C. Following blocking, samples were incubated with mouse anti-CD45:Alexa Fluor 700 (1:10, Cat. Nr. MCA43A700, Biorad) and mouse anti-RP-1:Alexa Fluor 647 (1:20, Cat. Nr. 550000, BD Pharmingen) for 30 min at 4°C in the dark. After incubation, cells were washed and fixed in 4 % PFA for 15 min at 4°C in the dark, followed by washing and resuspension in 100 µL FCSB. To obtain absolute cell counts, 100 µL of AccuCheck Counting Beads (Thermo Fisher Scientific) were added to each sample by reverse pipetting. Stained cells were acquired using a BD LSRFortessa X-20 flow cytometer equipped with 5 lasers (488nm, 535nm, 633nm, 405nm, 355nm). A total of 10000 events were collected per sample. Compensation was applied using the AbC Anti-Mouse Bead Kit (Thermo Fisher Scientific), and fluorescence-minus-one (FMO) controls were used to set gating thresholds. Flow cytometry data were analyzed using FlowJo (version 10.7.1). Absolute cell counts were calculated according to the manufacturer’s instructions for AccuCheck Counting Beads and normalized to total vegetation weight.

### IL-1β cleaving assay

*E. faecalis* overnight cultures of OG1RF WT and mutants were diluted 1:10 in 200 μl fresh BHI supplemented with human pro-IL-1beta (100 ng/mL; Cat. Nr. 10139-H07E, Sino Biological) and incubated for 2, 4, 6 and 24 h at 37°C, static conditions. Media was additionally supplemented with 25 µg/ml erythromycin when *ΔgelE* pTCV-Ptet and *ΔgelE* pTCV-Ptet-*gelE* were tested. After incubation, bacteria were pelleted, and supernatants were stored at -20°C until separation with SDS-PAGE and western blotting. Sterile BHI media supplemented with human pro-IL-1beta served as a negative control. For mass spectrometry analysis, supernatants harvested at 0, 6 and 18 h were separated with SDS-PAGE and stained with Coomassie Blue. Protein bands of interest were excised and stored in miliQ water. Peptide mass spectrometry and analysis was performed at the Taplin Mass Spectrometry Facility, Harvard Medical School, Boston, Massachusetts, USA.

### Mammalian cell culture

HEK-Blue IL1R cells (invivogen), which is a HEK 293 reporter cells for human and murine IL-1α & IL-1β cytokines, were cultured in DMEM (Gibco) supplemented with 10% (v/v) heat-inactivated fetal bovine serum, 100 U/mL penicillin, 100 µg/mL streptomycin, and 1x HEK-Blue selection (invivogen) at 37°C in a humidified incubator with 5% CO₂. Cells were passaged when they reached 80% confluency. For passaging or harvesting cells for assays, cells were washed and incubated in PBS for 2 min at 37°C, followed by vigorous pipetting of PBS over the cell monolayer to detach the cells.

### IL-1 signaling assay

*E. faecalis* overnight cultures of OG1RF WT and mutants were diluted 1:10 in 1 mL fresh BHI supplemented with human pro-IL-1beta (100 pg/mL) and incubated for 6 h at 37°C, static conditions. After incubation, bacteria were pelleted, and supernatants were filter-sterilized. 20 µl of bacterial supernatant filtrate were added to 50,000 HEK-Blue IL-1R cells (invivogen) in 180 µl DMEM supplemented with 10% FBS and 100 U/mL Penicillin-Streptomycin in a 96- well plate. As negative controls, sterile BHI, DMEM and DMEM supplemented with human pro-IL-1beta (100 pg/mL) were used instead of bacterial supernatants. DMEM supplemented with human mature IL-1β (100 pg/mL) served as a positive control. Samples were incubated for 18 h at 37°C. The following day, 20 µl of HEK-Blue IL-1R cell supernatants were transferred to 180 µl of QUANTI-Blue Solution (invivogen) in a 96-well plate and incubated at 37°C for 1 h, followed by measuring absorbance at 620 nm in a plate reader (TECAN).

### PCR screening of clinical isolates

The presence of *fsrA* was detected by PCR amplification *in vitro* (ENVALVE cohort) or using BLAST (Pittsburgh cohort). Single colonies grown overnight on COS agar at 37°C were resuspended in DreamTaq Hot Start Green PCR Master Mix (Cat. Nr. K9021, Thermo Fisher Scientific) with primer 2 and 3 (**Table S4**) at a concentration of 1.6 µM. The PCR program consisted of an initial denaturation step at 94°C for 2 min, followed by 25 cycles of amplification (denaturation at 94°C for 30 s, annealing at 54.1°C for 30 s, and extension at 72°C for 1 min) and then a final extension at 72°C for 6 min. PCR products were analyzed by electrophoresis on a 1 % agarose gel and stained by GelRed. Amplified fragments of *fsrA* corresponded to 740 bp. OG1RF WT and Δ*fsr* were included as positive and negative control respectively.

### Crystal violet assay

Bacteria grown overnight in BHI were washed and diluted 1:100 in fresh BHI. 100 µL of each strain tested was transferred in triplicates in a tissue culture-treated or non-treated polystyrene 96-well plate and incubated for 24 h at 37°C. At the incubation endpoint, wells were washed with PBS to remove non-attached bacteria and then 95 % ethanol was added for 15 min at room temperature to fix the biofilm. Ethanol was removed and wells were washed twice in PBS, before staining biofilm with 0.1% crystal violet for 10 min. Wells were washed 3 times with PBS, followed by addition of 95 % ethanol for 10 min to dissolve crystal violet and mixing. Absorbance at 540 nm was measured in each well using a spectrophotometer.

### Microbroth dilution assay

Bacteria grown overnight in BHI were washed and resuspended in cation-adjusted Mueller-Hinton Broth (MHB). Bacterial concentration was adjusted to 1 x 10^6^ CFU/mL in MHB and incubated with a 2-fold serial dilution of ampicillin (0.25 – 16 µg/mL) and gentamicin (2 – 128 µg/mL) in a 96-well plate for 18 h at 37°C without shaking. Wells without antibiotics served as controls. Growth was visually assessed at the incubation endpoint.

### Statistical analysis

Statistical analysis for *in vivo* and *in vitro* assays was performed using GraphPad Prism 6 (version 6.07). One-way ANOVA with Tukey’s multiple comparison test was applied to identify significant differences among the mean values of gene expression (Log2FC) across different shear stress conditions, and mean absorbance values across different supernatants tested on HEK-Blue IL1R cells. A two-tailed t-test was applied to identify significant differences in the mean ΔCt between bacterial inoculum and 72-h vegetations, mean biofilm, and microcolony area between 2 experimental animal groups, and mean protein band density between 2 samples. Mann-Whitney U test was applied to compare the median CFU, weight, and cell count values between two experimental animal groups. Two-way ANOVA was applied to identify significant differences in the mean absorbance values among of WT and *lrgAB* mutant strains grown in 2 different media. Statistical data analysis of the clinical data was conducted using R (version 2024.04.2+764, R Core Team, Vienna, Austria). Given the uneven distribution of *fsrA* presence between the 2 cohorts, we aimed to confirm the absence of a correlation between bacteraemia duration and cohort depending on gene presence. To this end, a gamma generalised linear model was applied to examine the effect of cohort within the *fsrA*- present and *fsrA*-absent groups separately. To evaluate the relationship between gene presence and various clinical and demographic variables a contingency table analysis was performed. Contingency tables were generated to summarise the association between gene presence and each of the selected variables, as well as gene presence and a cumulative score. The scoring system was established with a maximum of 7 points describing the highest disease severity. Patients received 2 points for the fulfilment of the modified Duke Criteria ^70^ to classify the disease as “definite IE”, or 1 point when classified as “possible IE”. Further to represent the severity of the individual clinical course, one point for each of the following item was received: In-hospital mortality, necessity of treatment on the ICU, as well as ICU length of stay equalling 14 or more days. As a representation of the local extent of the infection one point for each of the following item was received: echocardiographic evidence of vegetation as well as performance of valve replacement or device extraction, such as infected pacemaker electrodes, due to IE. Fisher’s exact test was used to assess the statistical significance of these associations. Heatmaps were created for each contingency table using ggplot2 to visualise the frequency of combinations between *fsrA* presence and the selected variable categories, with colour gradients indicating frequency intensity. To compare the baseline characteristics of both studies, descriptive statistics were generated using the gtsummary package to summarise categorical and continuous variables by the presence or absence of *fsrA* (**Supplementary File 3**). Categorical variables were summarised as counts and percentages, while continuous variables were reported as medians with interquartile ranges (IQR). Comparisons between groups were performed using the appropriate statistical tests: Chi-square tests or Fisher’s exact tests for categorical variables and Wilcoxon rank-sum tests for continuous variables. Subgroup analyses were performed for the Pittsburgh and Zurich cohorts, and results were compared across these cohorts. A combined summary for both cohorts was also generated.

## RESOURCE AVAILABILITY

### Lead contact

Further information and requests for resources and reagents should be directed to and will be fulfilled by the lead contact, Kimberly Ann Kline.

### Materials availability

Bacterial mutant strains generated, and clinical isolates used in this study will be made available on request, with reasonable compensation by requestor for shipping costs. We may require a completed materials transfer agreement if there is potential for commercial application.

### Data and code availability

Transcriptomic data have been deposited in NCBI under the bioproject accession numbers PRJNA1219810 and PRJNA1219807 and are publicly available as of the date of publication. Original western blot and microscopy images in this paper will be shared by the lead contact upon request. This paper does not generate original codes. Any additional information required to reanalyze the data reported in this paper is available from the lead contact upon request.

## ACKNOWLEDGEMENTS

Funding for this work was provided by the Singapore National Research Foundation and Ministry of Education Singapore under its Research Centre of Excellence Program (SCELSE). Funding by Singapore Ministry of Education (MOE2019-T2-2-089) was awarded to K.A.K, and the National Medical Research Council Open Fund (MOH-000645) to K.P. and K.A.K. This study was also supported by the Wallenberg Foundation Postdoctoral Fellowship at Nanyang Technological University Singapore to H.A., SCELSE Seed Funding (SF-05) to H.A., Open Fund – Young Individual Research Grant by the National Medical Research Council Singapore (MOH- 000939) to H.A., the University Of Zurich CRPP ‘Personalized Medicine Of Persisting Bacterial Infections Aiming to Optimize Treatment and Outcome’ to A.S.Z., B.H. and S.D.B. and by the Schweizerischer Nationalsfonds (SNF) (grant 310030_204343 to A.S.Z., grant 211422 to S.D.B, grant 32003B_219351 to B.H, and grant 310030_219227 to K.A.K.). This work was also supported by National Institutes of Health grant R21AI164018 to D.V.T., and by the Department of Medicine at the University of Pittsburgh School of Medicine. Y.L was also supported by the Pitt-Tsinghua Partnership Program. The funders had no role in study design, data collection and analysis, decision to publish, or preparation of the manuscript.

We thank Gary Dunny and Jennifer Dale for providing the *E. faecalis* transposon mutants used in this study. We are grateful for insightful discussions about this project with Alex Persat (EPFL), Carey Nadell (Dartmouth), and Irina Afonina (SMART). We also thank our colleagues at the SCELSE Sequencing Facility for performing library preparation and RNA sequencing. We also thank the instructors René Remie and Irene Cuesta at the René Remie Surgical Skills Centre for teaching us the surgical techniques required for the IE animal model. We are also grateful to Antonin André (University of Geneva) for critical feedback on the manuscript.

## AUTHORS CONTRIBUTIONS

Conceptualization: H.A., K.A.K.

Methodology: H.A., V.S., W.I.S., Y.L., S.D.B., S.L.W., D.V.T., A.S.Z., K.A.K.

Formal Analysis: H.A., V.S., W.I.S., Y.L., S.D.B., A.Z., D.V.T., A.S.Z., K.A.K.

Investigation: H.A., V.S., W.I.S., Y.L., R.J.W.T., K.K.F.N., C.J.Y.N., S.M.R., F.R.T., R.A.G.D.S., C.C.W., C.M., J.J.W.

Resources: H.A., K.P., S.D.B., S.L.W., D.V.T., A.S.Z., K.A.K.

Writing – Original Draft: H.A.

Writing-Review and editing: V.S., W.I.S., Y.L., R.J.W.T., K.K.F.N., C.J.Y.N., S.M.R., F.R.T.,

R.A.G.D.S., C.C.W., C.M., J.J.W., K.P., B.H., S.D.B., S.L.W., D.V.T., A.S.Z., K.A.K.

Visualization: H.A., V.S. Supervision: H.A., S.L.W., K.A.K.

Funding Acquisition: H.A., K.P., B.H., S.D.B., S.L.W., D.V.T., A.S.Z, K.A.K.

## SUPPLEMENTAL INFORMATION

**Supplementary figures and tables.** Document containing Supplementary Figures S1-S6, and Supplementary Tables S1-S4

**Supplementary File 1.** Differential gene expression of *E. faecalis* under fluid flow compared to static conditions

**Supplementary File 2.** Differential gene expression of *E. faecalis* Δ*fsr* compared to WT in vegetation at 72 hpi

**Supplementary File 3.** Baseline characteristics of the study cohorts, by fsrA presence

