## Supplementary figures and tables for "Fsr quorum sensing system restricts biofilm growth and activates inflammation in enterococcal infective endocarditis"

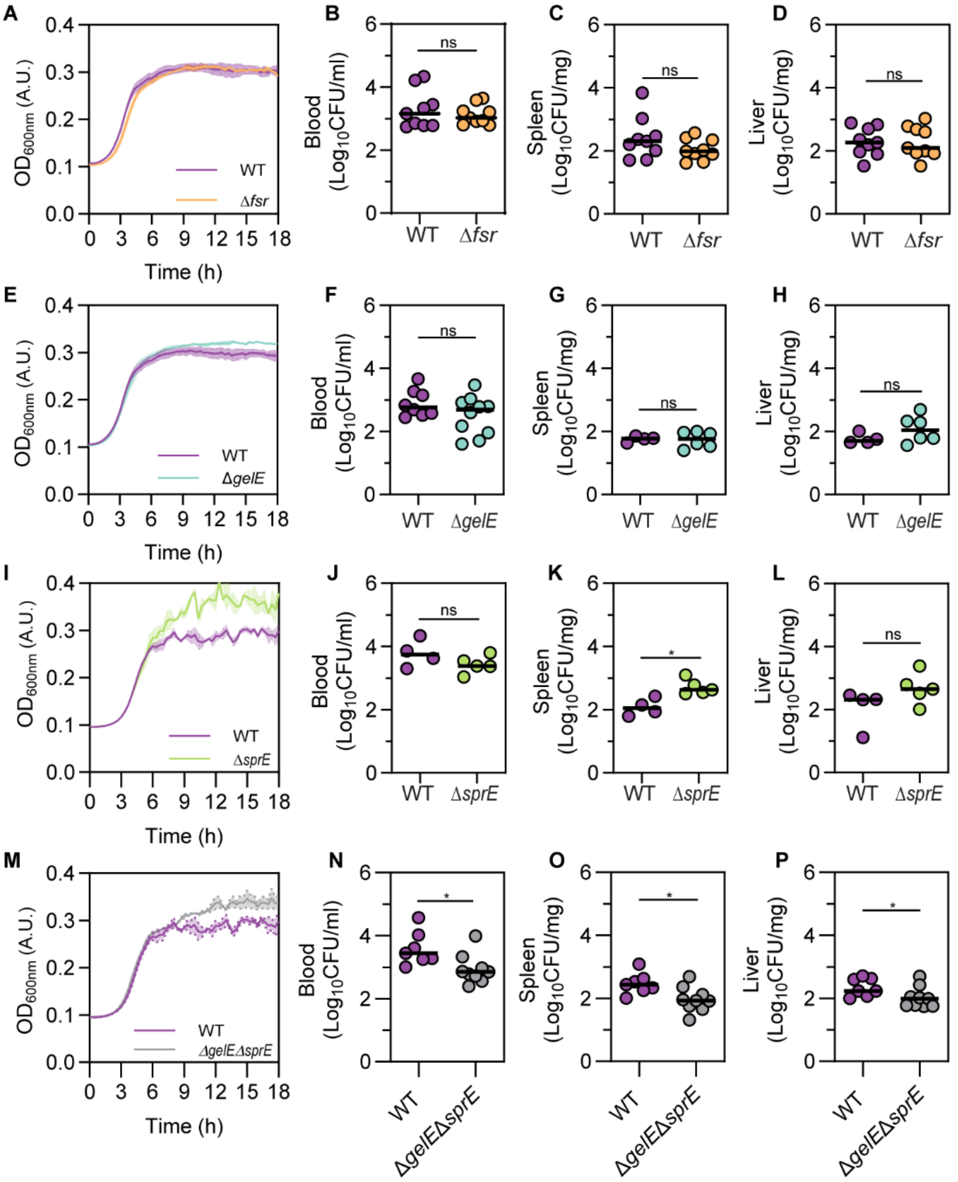


**Fig. S1. Growth and systemic spread of *E. faecalis* in the absence of Fsr QS system, *gelE*, and *sprE* in IE at 72 hpi. A, E, I, M.** Growth curves in BHIS at 37°C under aerobic conditions without shaking. Error = SEM represented as shaded region. N = 3 independent experiments for (A) and (E), N = 1 for (I) and (M). **B-D, F-H, J-L, N-P**. Median of blood, liver, and spleen CFU at 72 hpi. n = 9 animals per group from N = 2 is shown for (B-D), n = 8 - 10 animals per group from N = 2 for (F), n = 4 – 6 from N = 1 for (G-H), n = 4 – 5 from N = 1 for (J-L), n = 7 – 9 from N = 2 for (N-P). Statistical significance in median difference was assessed by applying a Mann-Whitney test; * = $p < 0.05$, ns = not significant.

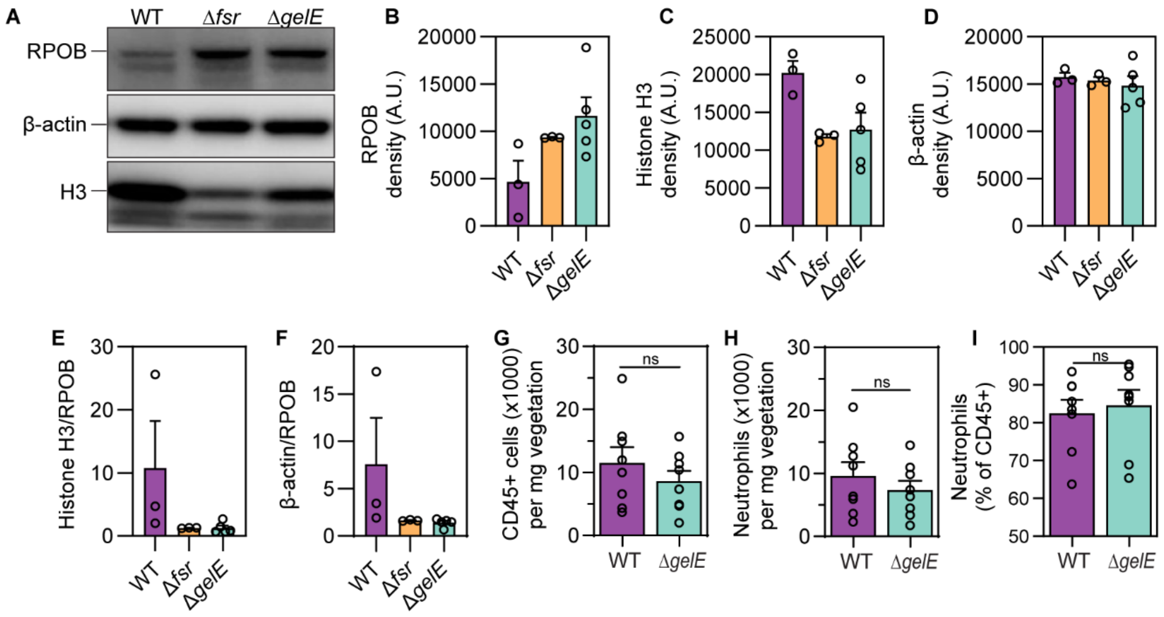


**Fig. S2**. **Increased bacterial load in Δ*fsr*- and Δ*gelE*-infected vegetations does not promote a further increase in host cell infiltration**. **A-D.** *E. faecalis* RPOB, and rat histone H3 and β-actin detection with western blotting in lysates from WT, Δ*fsr*-, and Δ*gelE*-infected vegetations harvested at 72 hpi (A), and protein band density quantification with ImageJ (B-D). RPOB levels (B) corresponding to bacterial abundance are higher in Δ*fsr*- and Δ*gelE*-infected vegetations compared to WT. Histone H3 levels (C) corresponding to nucleated host cell abundance and β-actin levels (D) corresponding to both nucleated host cells and anucleate platelets are similar across all vegetations. Representative bands are shown in A. Samples were harvested from n = 3-5 animals per group from N = 1 independent experiment. Error = SEM. **E-F.** Histone H3 and β-actin levels normalized by RPOB levels, reflecting the relative abundance of host cells to bacteria. Analysis is based on results from (A-D). **G-I.** Leukocyte (CD45+) (G) and neutrophil (H) absolute quantification, and neutrophil relative quantification (I) at 72 hpi vegetations using flow cytometry. Neutrophils (% of CD45+ cells) were determined based on the number of CD45+ RP-1+ events of the total CD45+ events. Mean with SEM is shown from n = 8 animals per group from N = 2 independent experiments. Statistical significance was assessed with a t-test; ns = not significant

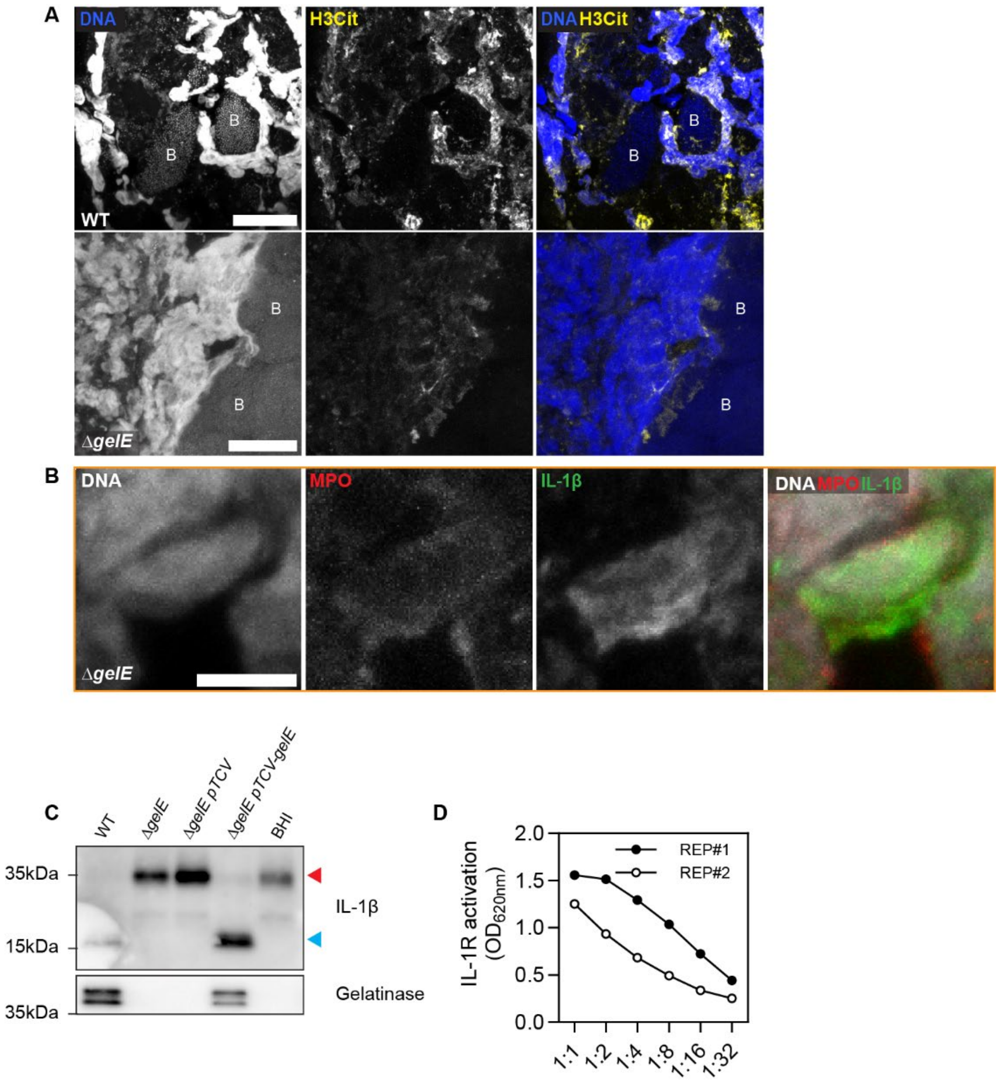


**Fig. S3.** ***E. faecalis*** **gelatinase cleaves and activates IL-1β. A.** Neutrophils undergoing NETosis at the interface with biofilm (B), evidenced by their decondensed nuclei colocalizing with citrullinated histone H3 (H3Cit) in WT- and Δ*gelE*-infected vegetations at 72 hpi. Representative Z-projection captured with LSCM from n =3 animals per group from N = 1 is shown. DNA is stained with DAPI. Scale = 20 µm. **B.** Inset (orange) from Fig. 5A (Δ*gelE* panel) showing IL-1β colocalizing with a decondensed nucleus of an incoming neutrophil. Colocalization with myeloperoxidase (MPO) indicates that the nucleus belongs to a neutrophil. Scale = 5 µm. **C.** Complementation of Δ*gelE* with pTCV-Ptet-*gelE* (Δ*gelE* pTCV-*gelE*) restored secretion of gelatinase in the supernatant and cleaving of pro-IL-1β (blue arrowhead) to a 17 kDa fragment (red arrowhead). **D.** Activation of HEK-Blue IL-1R reporter cells in the presence of 2-fold serial dilutions of supernatants harvested from OG1RF WT incubated with pro-IL-1β for 18 h. N = 2.

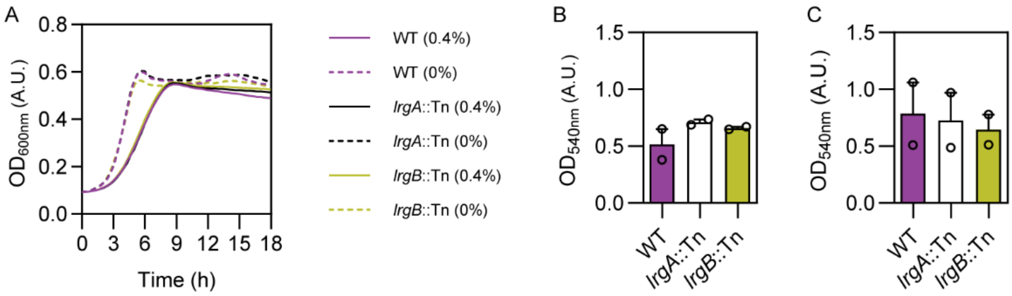


**Fig. S4. *lrgAB* is not involved in bacterial sensitivity to Triton X-100 and biofilm formation. A.** Growth curves of *E. faecalis* strains in BHI with 0.4 or 0 % Triton X-100 at 37°C under aerobic conditions without shaking. Mean of 3 technical replicates from N = 1 is shown. **B-C.** Biofilm formation at 24 h on tissue culture-treated (B) and uncoated polystyrene (C) 96-well plates assessed with the crystal violet assay. Mean of N = 2 is shown.

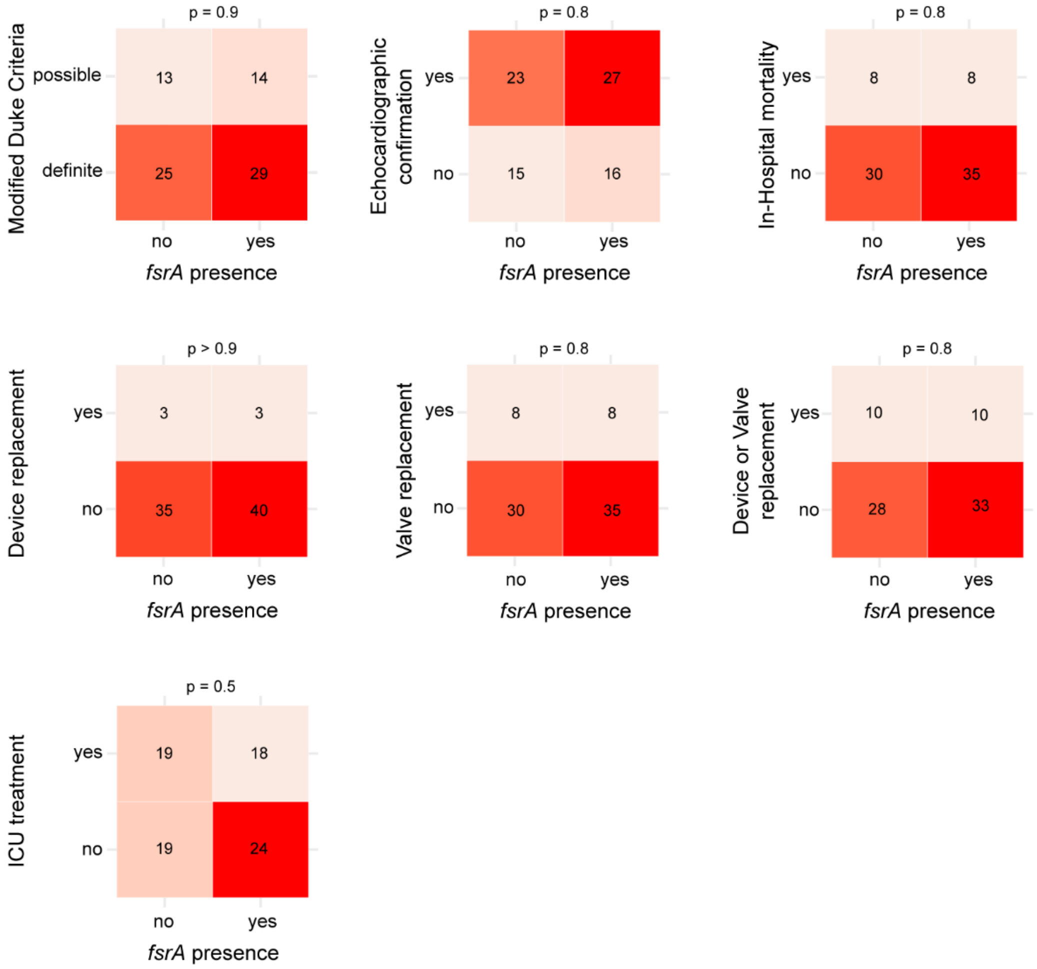


**Fig. S5. Relationship between *fsr* presence and clinical parameters of IE.** Contingency tables of the association of *fsrA* presence with different clinical parameters are shown. Statistical significance was assessed by Fisher's exact test.

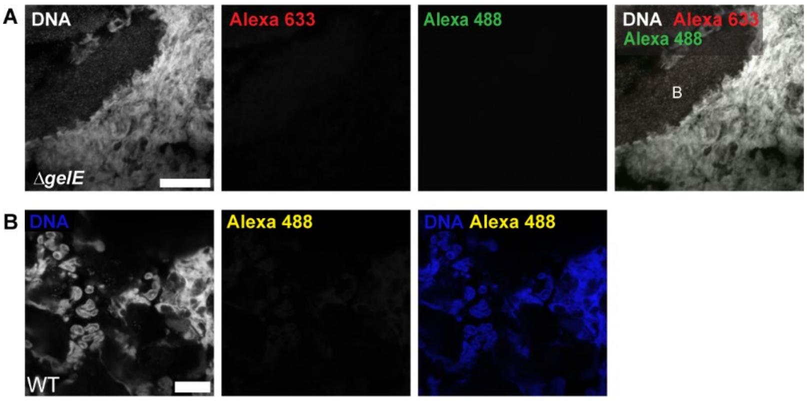


**Fig. S6. Secondary antibody only controls. A.** Secondary antibody only control performed on a tissue section consecutive to the section shown in Fig. 5A (Δ*gelE* panel). Z-projection of images captured with LSCM is shown, stained for DNA, goat anti-rabbit IgG Alexa 488, and goat anti-mouse IgG1 Alexa 633. B = biofilm, scale = 20 µm. **B.** Z-projections of images captured from WT-infected vegetations at 72 hpi captured with LSCM, stained for DNA and goat anti-rabbit IgG Alexa 488. Representative image of neutrophils undergoing NETosis from N=3 is shown. Scale = 20 µm.

**Table S1. MIC of ampicillin for *E. faecalis* OG1RF WT, *lrgA*::Tn, and *lrgB*::Tn strains**

| | MIC (μg/ml) | | |
|---|---|---|---|
| | **Rep#1** | **Rep#2** | **Rep#3** |
| **WT** | 0.5 | 1 | 2 |
| ***lrgA*::Tn** | 0.5 | 1 | 2 |
| ***lrgB*::Tn** | 0.5 | 1 | 2 |

**Table S2. MIC of gentamicin for *E. faecalis* OG1RF WT and Δ*fsr* strains**

| | MIC (μg/ml) | | |
|---|---|---|---|
| | **Rep#1** | **Rep#2** | **Rep#3** |
| **WT** | 32 | 16 | 32 |
| **Δ*fsr*** | 32 | 16 | 32 |

**Table S3. *E. faecalis* strains and plasmids used in *in vitro* and *in vivo* assays**

| Strain | Description | Reference |
|---|---|---|
| OG1RF | *Enterococcus faecalis, Rif^R^, Fus^R^* | [1] |
| Δ*fsr* | OG1RF *fsrABDC* deletion | This work |
| Δ*gelE* | OG1RF *gelE* deletion | [2] |
| Δ*sprE* | OG1RF *sprE* deletion | This work |
| Δ*gelE*Δ*sprE* | OG1RF Δ*gelE*Δ*sprE* deletion | This work |
| Δ*gelE*::*gelE*$^{E352A}$ | OG1RF Δ*gelE*::*gelE*$^{E352A}$ (proteolytically inactive gelatinase) | This work |
| Δ*gelE* pTCV-$P_{tet}$ | *gelE* deletion mutant with empty expression vector | This work |
| Δ*gelE* pTCV-$P_{tet}$::*gelE* | *gelE* deletion mutant with constitutively expressed *gelE* | This work |
| *sprE*::Tn | OG1RF, Tn insertion at site 1586796, $Cm^R$ | [3] |
| *lrgA*::Tn | OG1RF, Tn insertion at site 2597437, $Cm^R$ | [3] |
| *lrgB*::Tn | OG1RF, Tn insertion at site 2597099, $Cm^R$ | [3] |
| pGCP213 | Gram-positive, temperature-sensitive shuttle vector for allelic replacement | [4] |
| pTCV-$P_{tet}$ | Low-copy shuttle vector with constitutive $P_{tet}$ promoter for expression; $Kan^R$, $Erm^R$ | [5,6] |
| pTCV-$P_{tet}$::*gelE* | Complementation plasmid for *gelE* expression under constitutive $P_{tet}$ promoter; $Kan^R$, $Erm^R$ | This work |

**Table S4. Oligonucleotides used in this study**

| Oligo | Sequence (5'-3') | Description |
|---|---|---|
| fsrA_F | AGCAACCTCAAATCCTGCCT | qPCR |
| fsrA_R | TACAAGTGGCACACCAGGAC | qPCR |
| Primer 2 | ATGAGTGAACAAATGGCTATTTA[1] | FW primer, *fsrA* detection in clinical isolates |
| Primer 3 | CTAAGTAAGAAATAGTGCCTTGA[1] | RV primer, *fsrA* detection in clinical isolates |
| fsrB_F | AGACCTTGGATGACGAGACCG | qPCR |
| fsrB_R | GGTATGCGCCACAAGGAACA | qPCR |
| fsrC_F | TGCACTGTTTTCAATCGCGT | qPCR |
| fsrC_R | ACCGCAAAGCAAGCAAAACT | qPCR |
| gelE_F | ACAAGATGGGCATCCCTCGA | qPCR |
| gelE_R | TCAAGCGCCATCACTAGCGA | qPCR |
| sprE _F | ATTGCGGTAGTGACTGTCGG | qPCR |
| sprE_R | CGACCATTGCGTGTGGTTTT | qPCR |
| entV_F | AGCTGCACAAAAGAAAGCCTG | qPCR |
| entV_R | TAGCCCACATTGAACTGCCC | qPCR |
| RS04585_F | ACTTGGATAGATTCAGGAGAACTTG | qPCR |
| RS04585_R | TGGGTATTGCATAGAACATGGC | qPCR |
| recA_F | GCGGCTGTTCCACCATTTCG | qPCR |
| recA_R | GTTGCATTGGGCGTAGGTGG | qPCR |
| dnaB_F | CGTGGTGAAGGAGAAGACGGT | qPCR |
| dnaB_R | TCCTCGGCGAAAGCGAAGAA | qPCR |
| fsrABDC_del_1 | ATTGCTTA<u>CTCGAG</u>TGGCGTGAC[2] | *fsrABDC* upstream-F for Δ*fsrABDC* mutant construction |
| fsrABDC_del_2 | TTCGTTAACAACTTTTTTGTTCACTCATCCCTTTCTC | *fsrABDC* upstream-R for Δ*fsrABDC* mutant construction |

| | | |
|---|---|---|
| fsrABDC_del_3 | GGATGAGTGAACAAAAAAGTTGTTAACGAATGAATTTG | *fsrABDC* downstream-F for $\Delta$*fsrABDC* mutant construction |
| fsrABDC_del_4 | CCGTTTGCTTTTCAAGCTTAAGATGCTT[2] | *fsrABDC* downstream-R for $\Delta$*fsrABDC* mutant construction |
| sprE-del-1 | GTTGACCGAAAAACAGGAATTCGAAATTTA[2] | *sprE* upstream-F for $\Delta$*sprE* mutant construction |
| sprE-del-2 | CACAGCGGATAAACGGATCATGCCACTCCTTATCC | *sprE* upstream-R for $\Delta$*sprE* mutant construction |
| sprE-del-3 | TAAGGAGTGGCATGATCCGTTTATCCGCTGTGCCAGC | *sprE* downstream-F for $\Delta$*sprE* mutant construction |
| sprE_del_4 & gelE_del_4 | AAAGAAAGAAAGCTTGTCACAGATAAAACG[2] | *sprE* downstream-R for $\Delta$*sprE* mutant construction |
| gelE_del_1 | GTGTCCAAGCCGAATTCGATTTTAG[2] | *gelE* upstream-F for $\Delta$*gelE* $\Delta$*sprE* mutant construction |
| gelE_del_2 | GCACAGCGGATAAACGTTCCAACAAAGATGCCTGT | *gelE* upstream-R for $\Delta$*gelE* $\Delta$*sprE* mutant construction |
| gelE_del_3 | TACAGGCATCTTTGTTGGAACGTTTATCCGCTGTGCCAGC | *gelE* downstream-*F for* $\Delta$*gelE* $\Delta$*sprE* mutant construction |
| gelE_ins_1 | gatgcatgctcgagcGAATTGAAAATGTTCGCTATCTC[3] | *gelE* upstream-F for *gelE* chromosomal insertion |
| gelE_ins_2 | GAATAAACTTGTTCTTC**C**GCGGC[4] | *gelE* upstream-R with silent mutation for *gelE* chromosomal insertion |
| gelE_ins_3 | GTAGCCGC**G**GAAGAACAAG[4] | *gelE* downstream-F with silent mutation for *gelE* chromosomal insertion |
| gelE_ins_4 | taccgagctcggatcGACGATCGTTTTGTTTGC[3] | *gelE* downstream-R for *gelE* chromosomal insertion |
| pGCP213_F | GCTCGAGCATGCATCTAGAGG | pGCP213 inverse PCR-F |
| pGCP213_R | GATCCGAGCTCGGTACCAAG | pGCP213 inverse PCR-R |
| gelE_E352A_SDM_F | CAGGTGCCTTGAATG**C**ATCTTATTCTG[4] | *gelE*(E352A) site directed mutagenesis-F |
| gelE_E352A_SDM_R | CAGAATAAGAT**G**CATTCAAGGCACC[4] | *gelE*(E352A) site-directed mutagenesis SDM-R |
| gelE_pTCV-Ptet_F | ATGCCTATGGGATCCGAGGATAAAGCAATACTTTTGTTGG | *gelE* insert-F for plasmid complementation |
| gelE_pTCV-Ptet_R | CTTGCATGCCTGCAGGAATTTTTTTCATTCATTGACCAG | *gelE* insert-R for plasmid complementation |

[1]Nakayama J, Kariyama R, Kumon H, 2002

[2]Underlined nucleotides indicate restriction site

[3]15bp overhang of homologous region for InFusion (Gibson assembly) in lowercase

[4]Point mutations for SDM annotated in bold
